# Targeted reduction of highly abundant transcripts with *pseudo-random* primers

**DOI:** 10.1101/027805

**Authors:** Ophélie Arnaud, Sachi Kato, Stéphane Poulain, Charles Plessy

## Abstract

Transcriptome studies based on quantitative sequencing estimate gene expression levels by measuring the abundance of target RNAs in libraries of sequence reads. The sequencing cost is proportional to the total number of sequenced reads. Therefore, in order to cover rare RNAs, considerable quantities of abundant and identical reads have to be sequenced. This major limitation can be lifted by strategies used to deplete the library from some of the most abundant sequences. However, these strategies involve either an extra handling of the input RNA sample, or the use of a large number of reverse-transcription primers (termed “not-so-random primers”), which are costly to synthetize and customize. Here, we demonstrate that with a precise selection of only 40 “pseudo-random” reverse-transcription primers, it is possible to decrease the rate of undesirable abundant sequences within a library without affecting the transcriptome diversity. “Pseudo-random” primers are simple to design, and therefore are a flexible tool for enriching transcriptome libraries in rare transcripts sequences.

## Methods summary

The precise selection and the use of pseudo-random primers allows for reducing the detection of undesirable sequences within libraries and so increase the effective depth of the sequencing. Our study also concludes that, instead of the 4096 random primers currently used, only 40 pseudo-random primers are enough.

## Introduction

In transcriptome studies using quantitative sequencing, highly abundant sequences within a library limit the coverage and increase the difficulty to detect transcripts of interest. For example, ribosomal RNAs (rRNA) can represent the majority of a sequence library, which means that most of the money spent on sequencing would be for reads that are irrelevant in the downstream analysis. For this reason, transcriptome analysis methods often include a step for removing rRNA.

At present, several methods exist to deplete rRNA, for example, by priming the cDNAs or enriching the mRNAs with poly-T oligonucleotides, by capturing and removing the rRNAs with hybridization probes and magnetic beads (Ribo-Zero kit) (1) or antibodies directed against DNA:RNA hybrids (GeneReadrRNA depletion kit) (2), by capturing first-strand cDNAs synthesized from capped transcripts (CAP Trapper) (3), by selectively degrading the 5’-phosphate RNAs (“Terminator” enzyme) (Epicentre), or by biasing the reverse-transcription primers against the rRNA sequences (4).

In this last method, termed “not-so-random primers” (NSR), the cDNAs are primed with a mixture of the 749 out of 4096 random hexamers that do not have a direct match with the human ribosomal RNAs, leading to a reduction of these sequences from 78% to 13% (4). The major drawback of this method is that the pool of primers is prepared by synthesizing each primer individually, which makes customization costly when adding a linker tail or changing the target for depletion (for instance hemoglobin) (5).

Here, we present a dramatic simplification of the not-so-random primers concept, which we term “pseudo-random primers” (PS). Following the initial observation of Mizuno et al. (1999) that the reverse-transcriptase tolerates even two mismatches at the priming site (6), we reasoned that a large number of not-so-random primer sequences are functionally redundant and that it would be possible to dramatically reduce their number, thus facilitating the development and testing of custom sets.

## Materials and methods

### Selection of PS primers

The 40 PS primers were selected to bind neither to the human rRNA nor to the linker sequence of the template-switching oligonucleotide used in our experiments (See supplemental information 1).

The 40 primers were individually synthetized (Invitrogen) with standard desalting purification grade, resuspended at 100 μM in ultra-pure water and mixed equimolarly.

### Selection of PS_Hb primers

The 33 PS_Hb primers were selected as described in supplemental material 1, by discarding hexamers sequences targeting human α-globin RNA and human β-globin RNA.

### Library preparation

NanoCAGE libraries were prepared according to Salimullah *et al.,* 2011 using 50 ng of total RNA extracted from HeLa and THP-1 cells lines (7). Technical triplicates of each nanoCAGE library were prepared from each RNA sample. Four libraries were made, to compare 1) Random hexamers (RanN6) versus PS primers, 2) RanN6, PS and 40 randomly picked RanN6 (40N6) primers, 3) RanN6, PS, 3 subsets of 20 PS and 1 subset of 10 PS primers, and 4) RanN6 versus PS_Hb primers. Thus, differences between RanN6 and PS primers, depleting rRNA and artifacts, were replicated in three independent experiments. Details of each nanoCAGE library are available in supplementary table 1.

### Data processing and analysis

The prepared libraries were individually paired-end sequenced on a MiSeq sequencer (Illumina) using the standard nanoCAGE sequencing primers (7). The sequencing data were analyzed using the workflow manager Moirai (8). Briefly, the reads were demultiplexed and trimmed to 15 bases with FASTX-Toolkit (http://hannonlab.cshl.edu/fastx_toolkit/). Then, the reads coming from rRNA or oligo-artifacts were removed with TagDust (version 1.13) (9) and the remaining reads were aligned to the human genome (hg19) with BWA (version 0.7) (10). Then, the non-proper paired reads and the PCR duplicates were filtered out with samtools (version 0.1.19) (11). Finally, the properly paired reads were clustered and analyzed as in Harbers *et al.,* 2013 (12) (the scripts used for the analysis are provided in supplemental materials 2).

## Results and discussion

We tested the pseudo-random primers concept using the nanoCAGE method for transcriptome profiling (13). In this method, 5’ adapters are introduced by template-switching oligonucleotides during the reverse transcription, where random primers are used to cover the non-polyadenlyated transcriptome. Thus, the undesirable sequences in nanoCAGE libraries come mostly from 2 sources: the ribosomal RNA and primers-primers artifacts. The rate of these undesired sequences becomes especially problematic when the quantity of starting material is lower than a nanogram. We therefore designed pseudo-random primers to reduce rRNA and primer-primers artifacts at the same time. Using scripts written in the R language (see Supplemental Information 1), we identified 40 hexamers that do not fully match with the human rRNA reference sequences, and do not show similarities up to 2 mismatches with the nanoCAGE linker sequence. We prepared a mixture of 40 reverse-transcription primers containing these hexamers (PS), to replace the standard reverse-transcription random primers (RanN6).

We tested the PS primers on three sets of triplicated libraries prepared from HeLa and THP-1 cell line total RNA. Using nanoCAGE libraries prepared with RanN6 primers as a control (Figure S1), we observed a significant decrease in reads matching to ribosomal RNA (Fig 1A). Primer artifacts were also reduced (Figure 1B), but the difference was only statistically significant for the THP-1 libraries: for one HeLa set of triplicates, there was no diminution, but the overall amount of artifacts was uniformly low, making it difficult to see any effect of the PS primers. To exclude the possibility that the observed effect of the PS primers comes only from the reduction of the hexamer diversity regardless of our selection, we included a control using 40 randomly picked hexamers (40N6). These libraries did not significantly deplete rRNA reads, but had an impact on primer artifacts. We explain this effect by the fact that only a few hexamers match to the linker sequences of the nanoCAGE primers, and therefore the 40N6 set was depleted by chance. Indeed, only 32% of them match the linkers with no or 1 mismatch (Figure S2). This confirms the efficiency of our precise selection of the PS primers to decrease the detection of the undesired sequences within nanoCAGE libraries.

**Figure 1:**
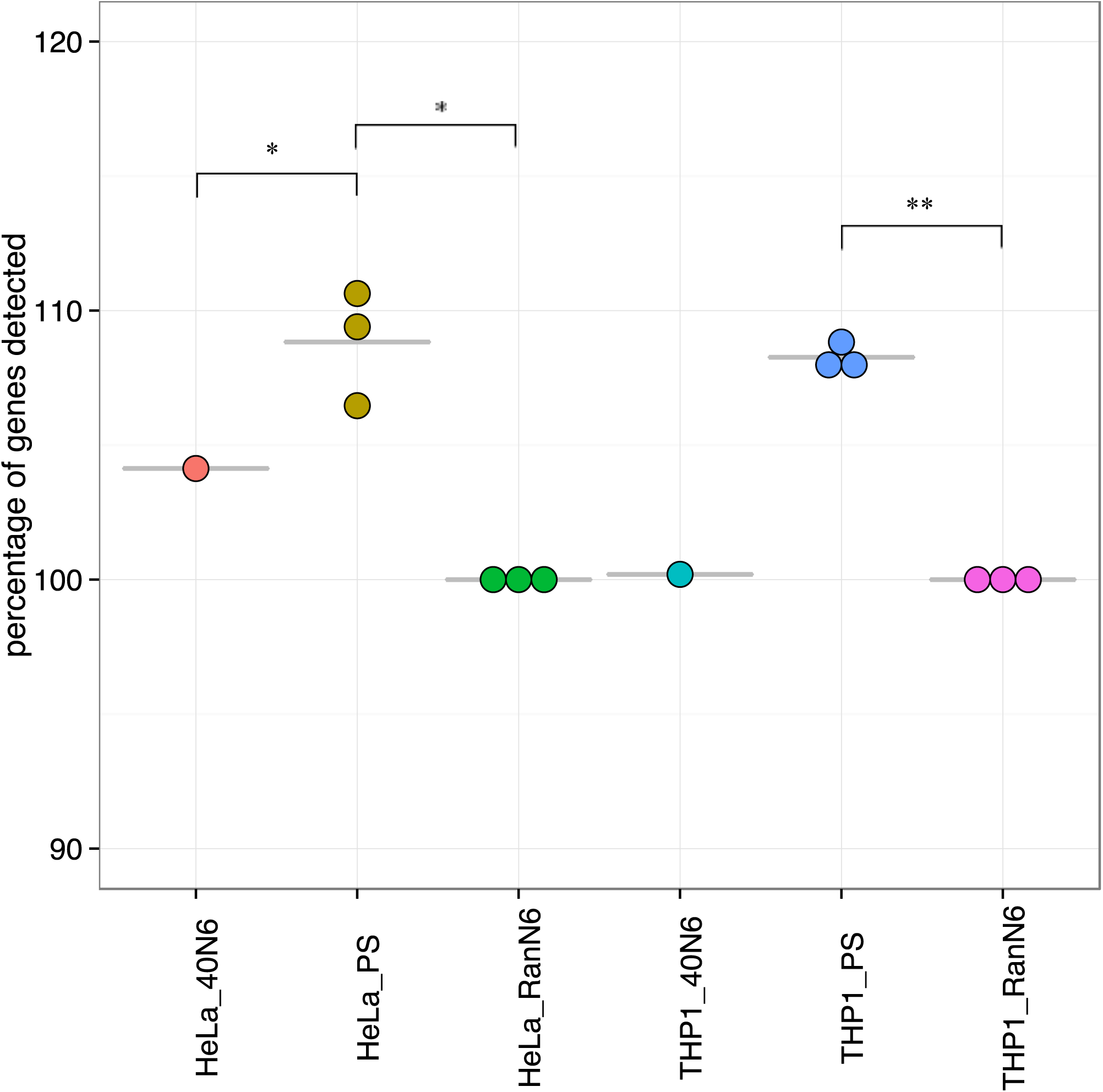
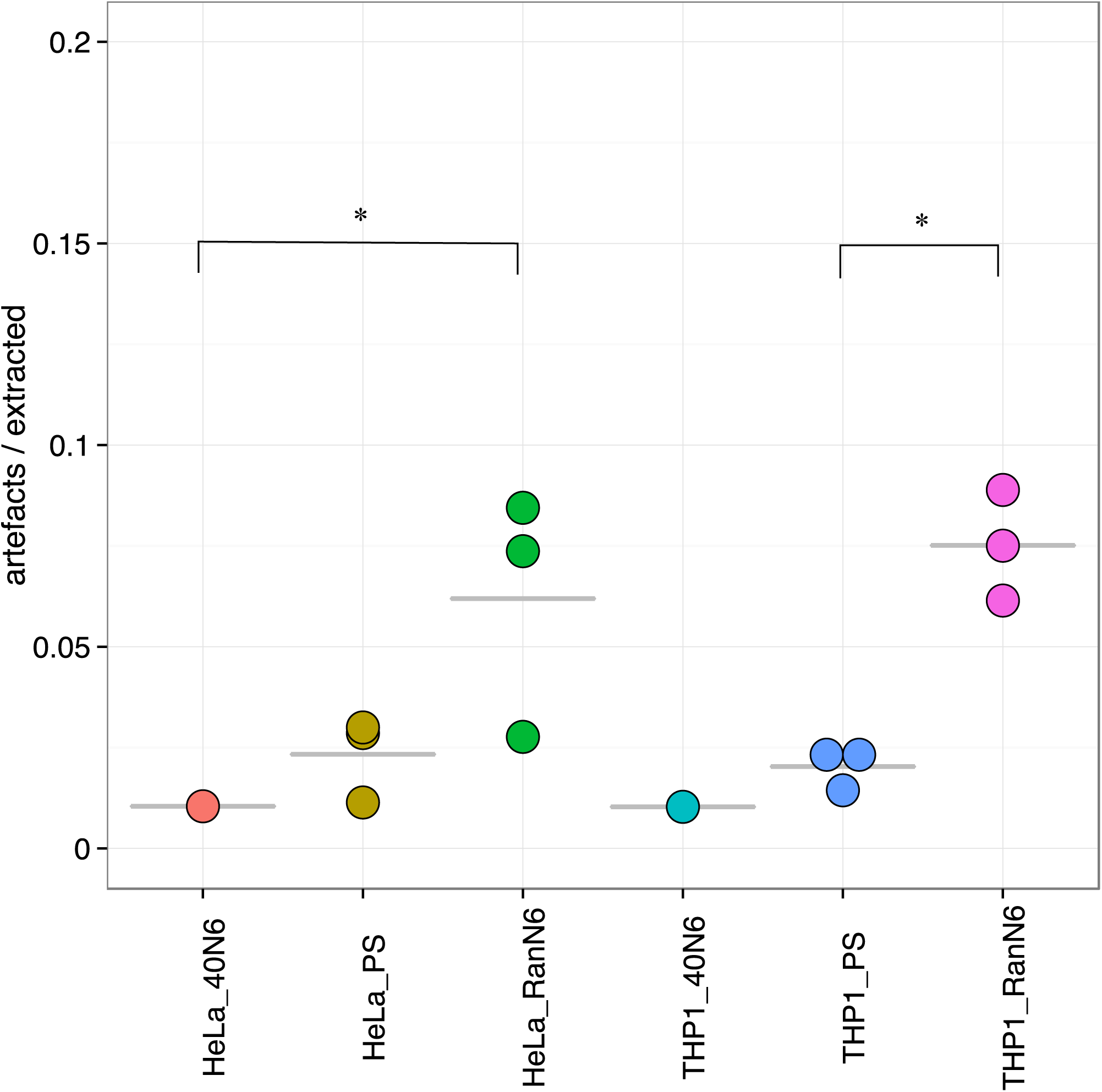
Depletion of ribosomal sequences and artifacts. Rate of ribosomal RNA (A) and artifacts (B) detected with the 40N6, PS or RanN6 primers sets. Each point corresponds to the mean of 3 technical replicates in the same experiment. Statistical test: t.test paired between the mean of PS and RanN6 data sets, non-paired with the raw value of 40N6 data set. * P-value<0.05; ** p-value<0.01; *** p-value<0.001.

We then verified that the two-fold reduction of the number of different hexamers did not impair genes detection. After normalizing the libraries to the same number of aligned reads (supplemental material 2), we detected between 3348 and 4235 genes per replicate (supplemental table 1). Not only the number of genes detected was not reduced with the use of only 40 primers, but also we detected significantly more genes with the PS primers than with the RanN6 primers, in both cell lines tested (Figure 2A). One simple explanation could be that PS primers that don’t bind to the ribosomal RNA are free to bind transcripts of interest, which would increase the likelihood of less abundant RNAs reverse-transcription. This is corroborated by the observation that libraries using the 40N6 primers, not selected against rRNA, do not allow for higher gene detection rate in comparison with the RanN6 primers. Importantly, because we normalized the number of aligned reads after filtering out the ones aligning on the rRNA, the effect of the PS primers can not be explained by the higher coverage at an equal number of raw reads. Altogether, our results show that the libraries prepared with PS primers cover more genes than the libraries prepared with RanN6 primers.

**Figure 2:**
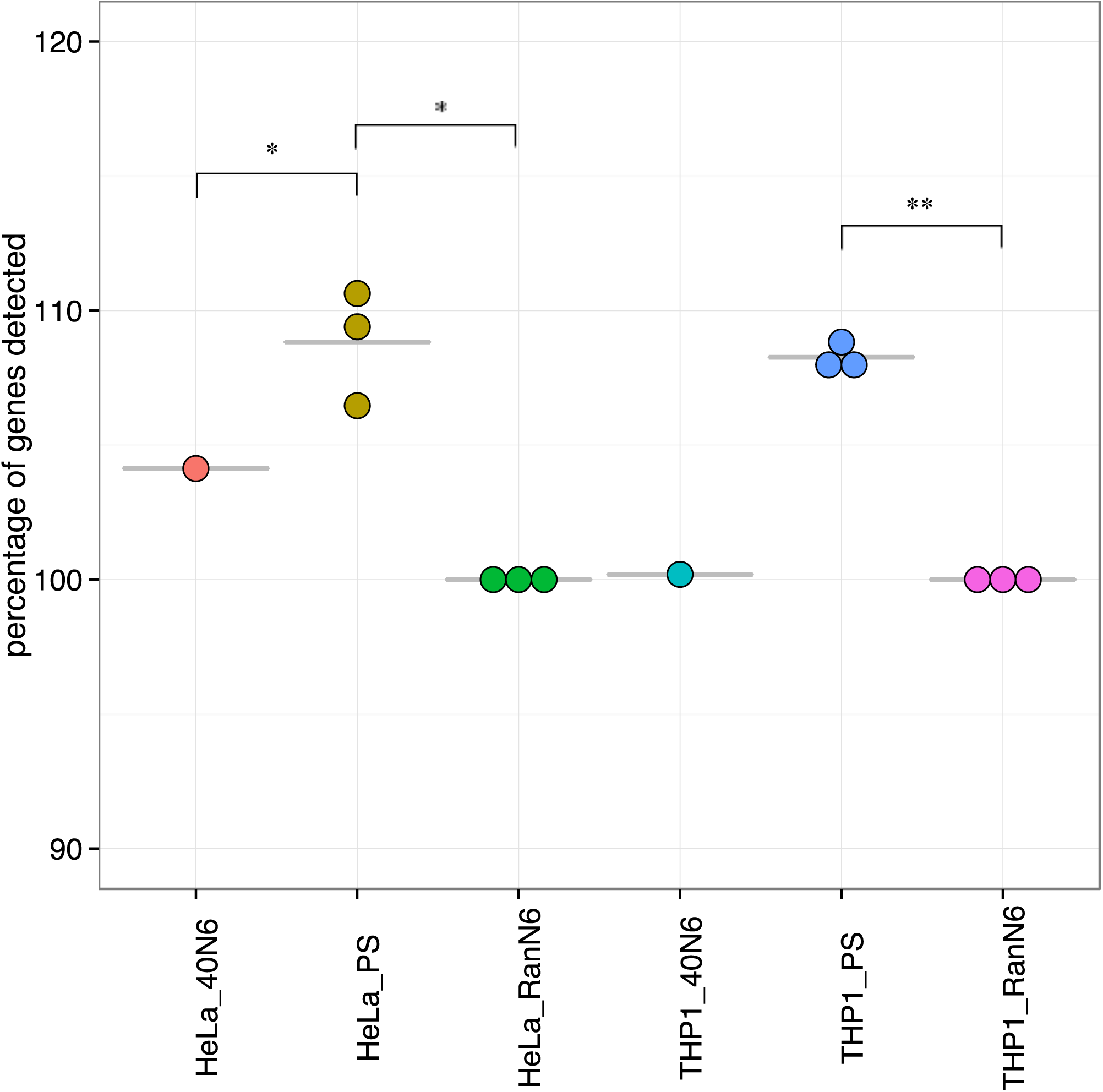
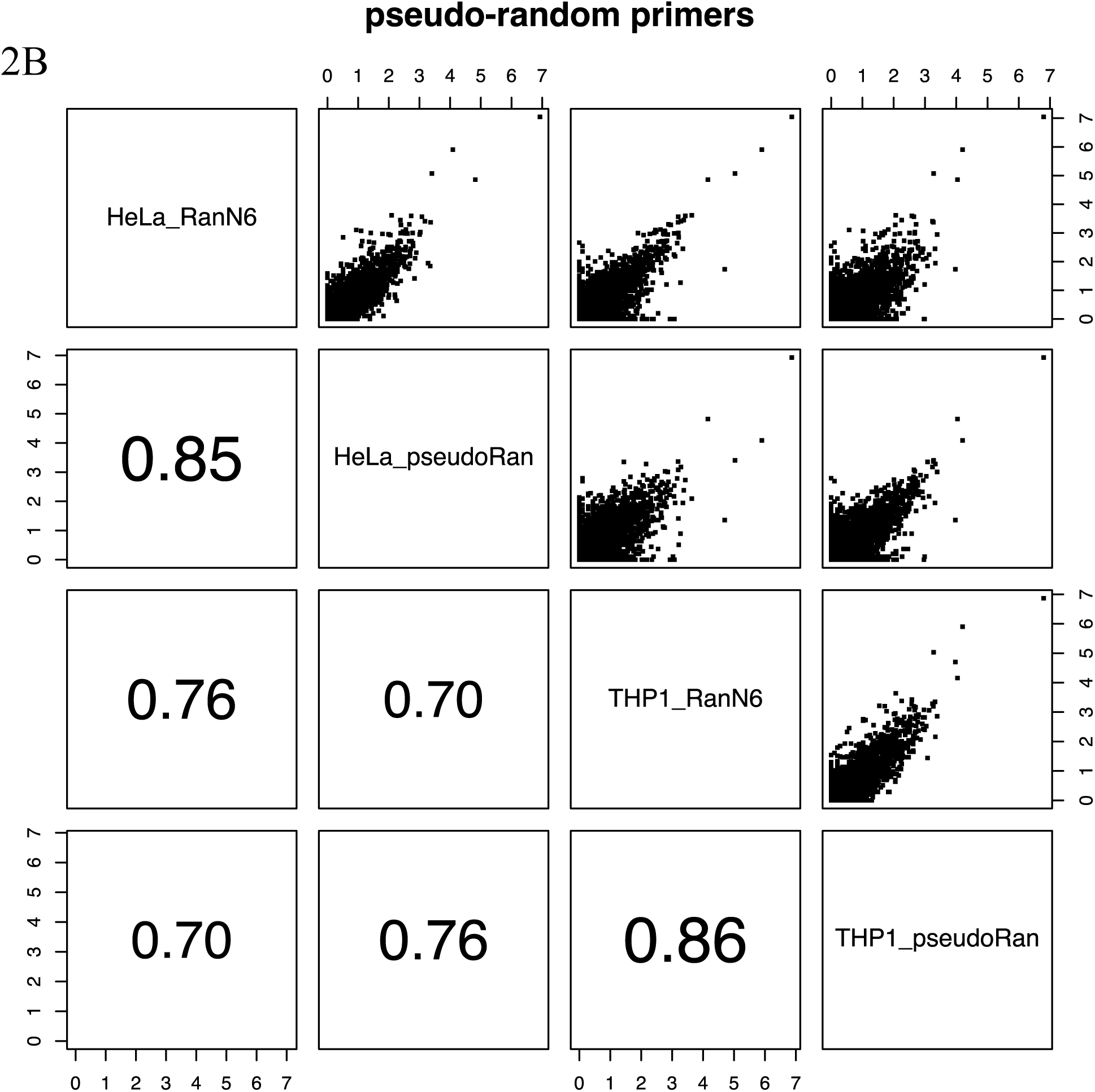
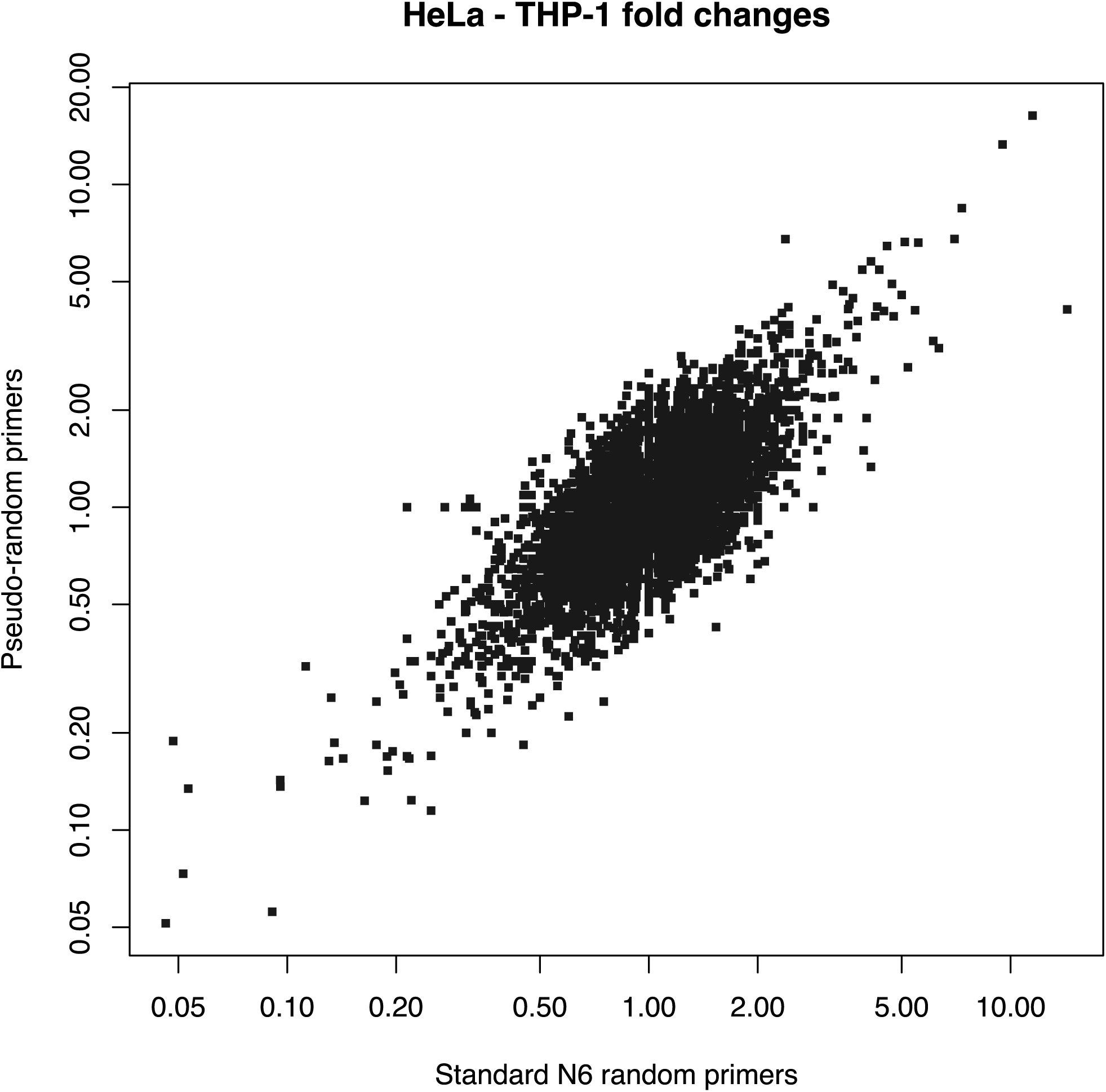
Coverage of transcriptome diversity. A. Percentage of genes detected with the 40N6 and PS primers compared to the RanN6 primers, set arbitrarily to 100 % in HeLa and THP-1 respectively. Each point corresponds to the mean of 3 technical replicates in the same experiment. The data were normalized by sub-sampling to 8700 tags per sample. Statistical test: t.test paired between the mean of RanN6 and PS data sets, non-paired with the raw value of 40N6 data set. * P-value<0.05; ** p-value<0.01; *** p-value<0.001 B. Pairwise comparison between the PS and RanN6 libraries from the 2 cell lines. Each plot is the mean of 3 experiments with 3 technical replicates. Upper part: expression plots where the reads are aligned to the reference gene model. Lower part: Pearson correlation of each pair. C. HeLa-THP1 fold change in gene expression.

To investigate the reliability of the expression values measured in PS-primed libraries, we compared our experiments pairwise after averaging the triplicates (supplemental material 2). Samples prepared from the same RNAs correlated better than samples prepared with the same RT primers set, but the correlation coefficients still suggested important differences induced by the change of primers (Fig 2B). Indeed, inspection of the pairwise plots shows that the most highly expressed genes deviate strongly from the diagonal when comparing the PS and RanN6 primers on the same RNA (Fig 2B). Given that the PS primers are strongly selected, this was expectable, and we reasoned that the bias should be systematic. To demonstrate that fact, we compared the fold change of expression levels between HeLa and THP-1 RNA in each set of primers, and showed that they were conserved (Fig 2C). Thus, libraries made with PS primers can be compared with libraries made with other RT primers by looking at fold changes with a common reference, like in transcriptome platform comparisons (14).

According to the good transcriptome coverage obtained with only 40 PS primers, we next wondered how many PS primers are required to conserve the same transcriptome diversity? The original number of 40 was set empirically from the matches on rRNA and nanoCAGE linkers, but the lower limit is unknown. We therefore prepared libraries with subsets of 20 or 10 PS primers (supplemental table 2). A similar number of genes (around 4000 genes per sample, supplemental table 1) could be detected in the libraries. We also observed a systematic bias in these libraries, but because they were made with subsets of the original PS primers, they all had a stronger similarity with each other than with RanN6 libraries (Fig 3). Thus, it appears possible to prepare whole-transcriptome libraries with as few as 10 pseudo-random primers.

**Figure 3:**
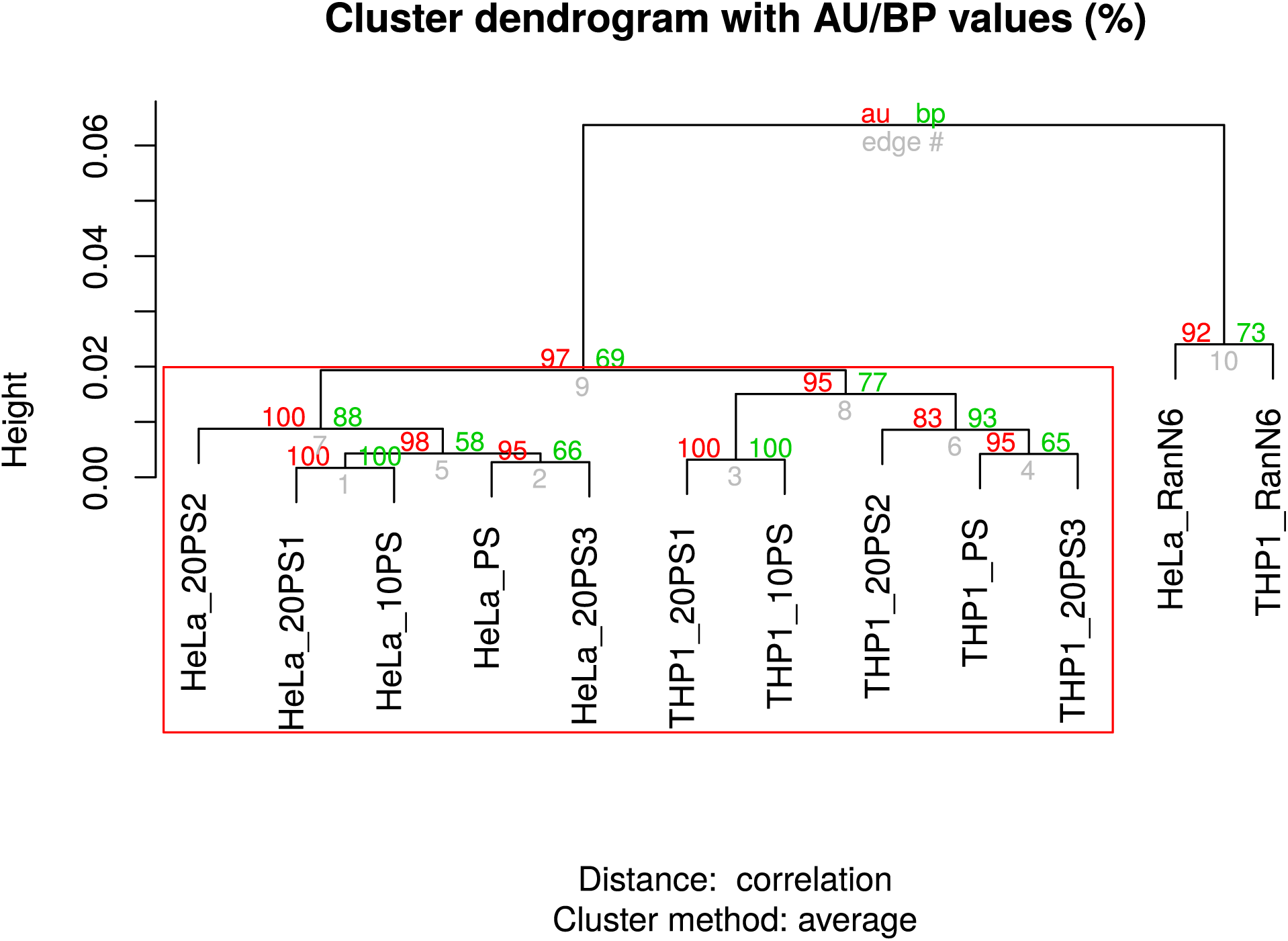
Transcriptome coverage with small number of primers. Hierarchical clustering of the detected genes (after normalization to 8700 tags per sample). The red value is the Approximately Unbiased (AU) p-value and the green value is the Bootstrap Probability (BP) value. The red box represent the cluster significantly established (AU p-value<0.05). All the samples were prepared in the same experiment (library NC_17).

Finally, we sought to demonstrate that the PS primers concept could be applied on other targets than the rRNAs. In total RNA extracted from blood, up to 60% of the transcripts come from hemoglobin genes, (15). Hence, we have selected 33 PS primers (PS_Hb) that did not match on hemoglobin sequences (with 2 or more mismatches) (supplemental material 1) and prepared nanoCAGE libraries with either these primers or standard RanN6 primers. The selection drastically reduced the number of tag per hemoglobin genes (Fig 4A), without reducing the number of detected genes (Fig 4B), thus demonstrating the possibility of designing PS primers against other targets.

**Figure 4:**
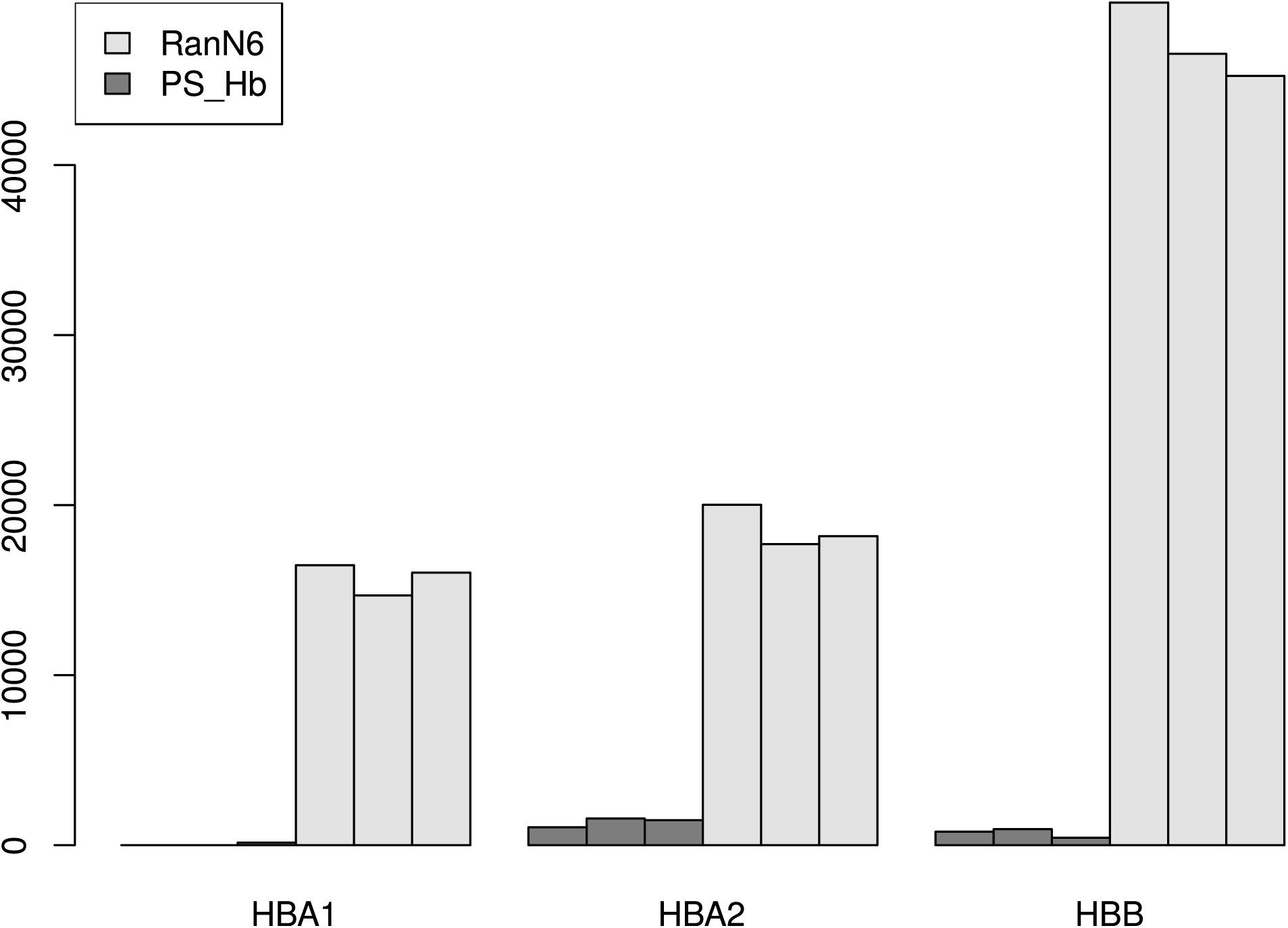
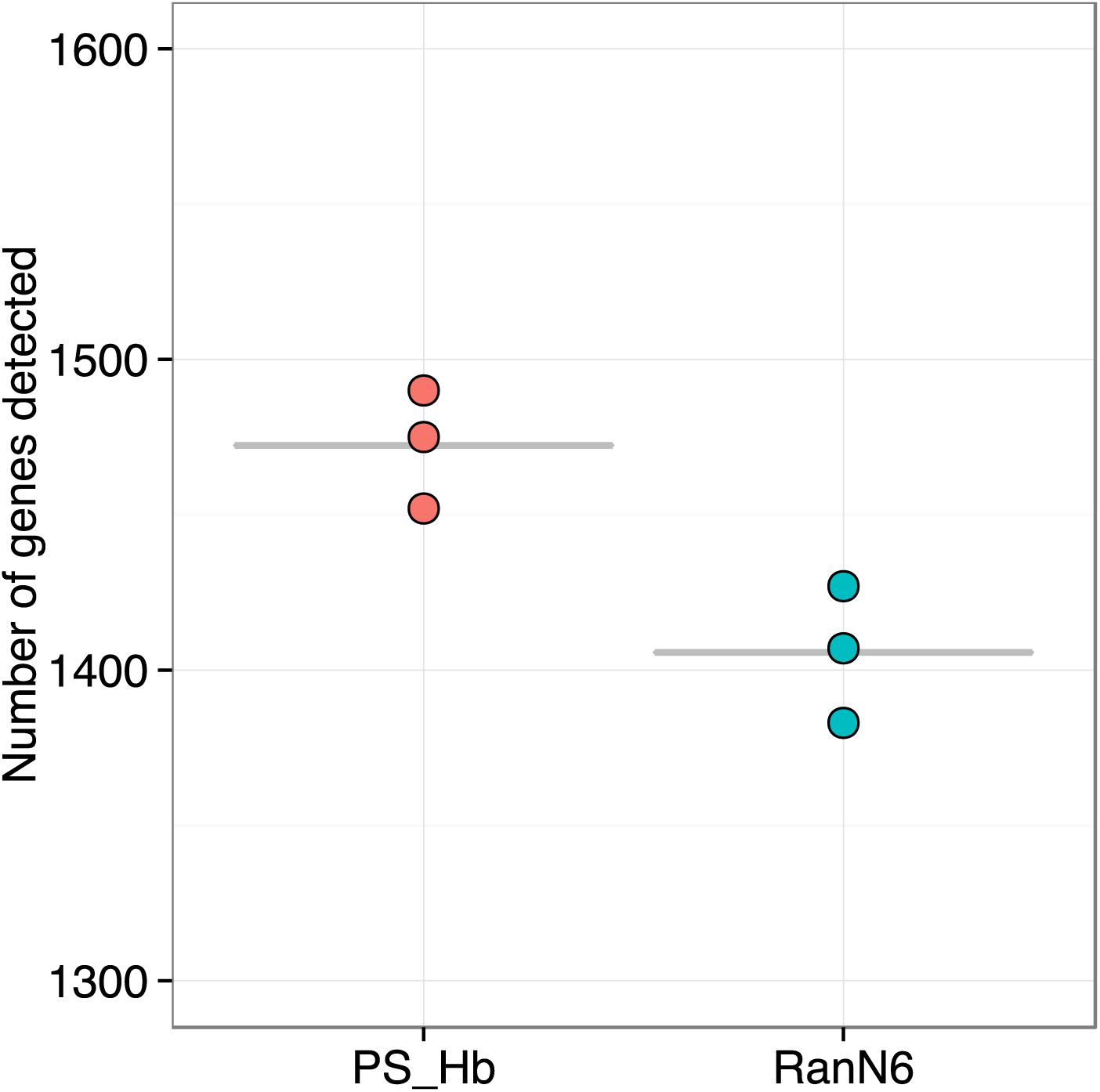
Targeted depletion of hemoglobin sequences. A. Measured expression levels (in counts per million) of hemoglobin genes with the PS_Hb and the RanN6 primers. Each bar represents a technical replicates of one experiment. B. Number of genes detected with the use of PS_Hb versus RanN6 primers. Each point corresponds to a technical replicate of the same experiment. The data were normalized to 3190 tags per sample.

In conclusion, despite several methods already exists to eliminate the sequences coming from ribosomal RNA in transcriptome studies, lots of them require an extra step in the protocol. Moreover, none of them is also able to eliminate, in a single step, multiple unrelated undesirable sequences. Here, we report that in transcriptome studies a drastic selection of the primers used during the reverse transcription is effective for eliminating specific sequences without reducing gene coverage. Moreover, our data supports the idea that the number of PS primers required is low, leading to a real cost-saving effect in the experiments. Finally, while tested here with the nanoCAGE protocol, this strategy is not limited to it and should be applicable to any kind of transcriptome studies.

## Authors contributions

CP conceived the project; OA, SK and CP designed the experiments; OA, SK and SP performed the experiments; OA and CP analyzed the data, OA and CP wrote the manuscript. All authors read and approved the final manuscript.

## Acknowledgements

The authors would like to thank Michiel de Hoon for the suggestion about using a subset of the pseudo-random primers; Roberto Simone who has suggested the name *pseudo-random primers* and Alistair Forrest and Yuri Ishizu for the gift of the human blood RNA sample.

This work was supported by a Research Grant from the Japanese Ministry of Education, Culture, Sports, Science and Technology (MEXT) to the RIKEN Center for Life Science Technologies, a Grant-in-Aid for Young Scientists A (number 25710018) and a research grant from the Mitsubishi Foundation (number 25142).

## Supplemental material

- Supplemental material 1: scripts and programs used for the primers selection
- Supplemental material 2: Scripts and programs used in the data analysis:

Link 1: general commands creating the files used in downstream analysis
Link 2: analysis of the first experiment, NCms10058
Link 3: analysis of the second experiment, NC12
Link 4: analysis of the third experiment, NC17
Link 5: common analysis of the three experiments
Link 6: Statistical analysis
Link 7: analysis of the fourth experiment regarding the RNA extracted from blood, NC22

- Supplemental material 3: Figure S2: Reads genomic features

Percentage of reads aligned to each feature of the genome (promoter, exon, intron, intergenique section, rDNA) and the artifacts. Each row is the average of the technical triplicates of the same library.

- Supplemental material 4: Figure S1: Maximal distance of the 40N6 primers with the template-switching primer

Number of mismatches between the hexamers of the 40N6 primers and the template switching primers.

- Supplemental material 5: Table S1: Summary table

Extensive summary for each sample tested. It includes the experiment name, the origin of the RNA, the barcode and index added, the primer set used and the sequencing results.

- Supplemental material 6: Table S2: sequences of the 20 PS and 10 PS primers sets.

Pseudo-random primers sequences composing the different sets of pseudo-random tested.

## Selection of Pseudo-random primers

### rRNA

Human:

hsu13369.fasta file produced with the command extractfeat -type rRNA U13369.gb. (from the EMBOSS package)

~~~
HSU13369_3657_5527 [rRNA] Human ribosomal DNA complete repeating unit.
HSU13369_6623_6779 [rRNA] Human ribosomal DNA complete repeating unit.
HSU13369_7935_12969 [rRNA] Human ribosomal DNA complete repeating unit.
~~~

### Mitochondrial

~~~
NC_012920_648_1601 [rRNA] Homo sapiens mitochondrion, complete genome.
NC_012920_1671_3229 [rRNA] Homo sapiens mitochondrion, complete genome.
~~~

### Combination

~~~
(cat nc_012920.fasta hsu13369.fasta  |  revseq -filter | grep -v ‘>‘  |  perl pe  chomp; echo) > ribo.txt
~~~

### R code

~~~
acgt <- c(‘A’, ‘C’, ‘G’, ‘T’)
LINKER <- ‘CCCTATAAGATCGGAAGAGCGGTTCGGAGACCTTCAGTTCGACTA’
BARCODES <- scan(‘barcodes.txt’, what=‘character’)
RIBO <- scan(‘ribo.txt’, what=‘character’) # See below in the wiki about the
  file ‘ribo.txt’.
hexamers <- apply(expand.grid(acgt, acgt, acgt, acgt, acgt, acgt), 1, paste,
collapse=‘‘)
hexamers <- data.frame(row.names=hexamers)
hexamers[,c(‘LINKER_0’, ‘LINKER_1’, ‘LINKER_2’, ‘LINKER_3’, ‘RIBO_0’, ‘RIBO_1
‘, ‘BARCODE’)] <- c(rep(FALSE, 7))
hexamers[names(unlist(sapply(rownames(hexamers), function(X) {agrep(X, LINKER
, 0, ignore.case=T)}))), “LINKER_0”] <- TRUE
hexamers[names(unlist(sapply(rownames(hexamers), function(X) {agrep(X, LINKER
, 1, ignore.case=T)}))), “LINKER_1”] <- TRUE
hexamers[names(unlist(sapply(rownames(hexamers), function(X) {agrep(X, LINKER
, 2, ignore.case=T)}))), “LINKER_2”] <- TRUE
hexamers[names(unlist(sapply(rownames(hexamers), function(X) {agrep(X, LINKER
, 3, ignore.case=T)}))), “LINKER_3”] <- TRUE
hexamers[names(unlist(sapply(rownames(hexamers), function(X) {agrep(X, RIBO
, 0, ignore.case=T)}))), “RIBO_0”] <- TRUE
hexamers[names(unlist(sapply(rownames(hexamers), function(X) {agrep(X, RIBO, 1, ignore.case=T)}))), “RIBO_1”] <- TRUE
hexamers[BARCODES, “BARCODE”] <- TRUE
~~~

~~~
summary(hexamers)
LINKER_0           LINKER_1           LINKER_2           LINKER_3           RIBO_0
    RIBO_1              BARCODE
  Mode :logical      Mode :logical      Mode :logical      Mode:logical      Mode :logical
       Mode:logical      Mode :logical
  FALSE:4056         FALSE:3082         FALSE:259         TRUE:4096         FALSE:719
      TRUE:4096         FALSE:4000
TRUE :40           TRUE :1014           TRUE :3837           NA’s:0           TRUE :3377
     NA’s:0           TRUE :96
NA’s :0            NA’s :0           NA’s :0                                 NA’s :0
~~~

~~~
with(hexamers, rownames(hexamers)[! (LINKER_2 | RIBO_0 | BARCODE)])
[1]    ”GCCAAA”    ”AGCAAA”    “AAACAA”    “ACACAA”    “TGCCAA”    “CAAACA”    “CACACA”    “TGCACA”
[9]    “GTCACA”    “TAGCCA”    “GTGGCA”    “TGTTTA”    “ATTTTA”    “CAAAAC”    “CACAAC”    “GCTAAC”
[17]    “AACCAC”    “CTACCC”    “TACCCC”    “CTAGCC”    “CTGGCC”    “TGTGCC”    “ATTGCC”    “CTACGC”
[25]    “TATGGC”    “TTGTGC”    “ACCACG”    “CACAGG”    “ACTGTG”    “TGCCAT”    “TGGCAT”    “GTGCAT”
[33]    “TTGTAT”    “ATTTAT”    “TTTTAT”    “TGGCGT”    “TGTTGT”    “ATTTGT”    “TTGCTT”    “TGTCTT”
~~~

## Selection PS_Hb

### Haemoglobin sequences

alpha globin mRNA: http://www.ncbi.nlm.nih.gov/nuccore/NM_000558 (http://www.ncbi.nlm.nih.gov/nuccore/NM_000558)

beta globin mRNA: http://www.ncbi.nlm.nih.gov/nuccore/NM_000518 (http://www.ncbi.nlm.nih.gov/nuccore/NM_000518)

The 2 fasta files are combined in 1 file named Hb.txt

### R Code

~~~
acgt <- c(‘A’, ‘C’, ‘G’, ‘T’)
Hb <- scan(‘Hb.txt’, what=‘character’)
hexamers <- apply(expand.grid(acgt, acgt, acgt, acgt, acgt, acgt), 1, paste, collapse=‘‘)
hexamers <- data.frame(row.names=hexamers)
hexamers[,c(‘Hb_0’, ‘Hb_1’, ‘Hb_2’)] <- c(rep(FALSE,3 ))
hexamers[names(unlist(sapply(rownames(hexamers), function(X) {agrep(X, Hb, 0, ignore.case=T)}))), “Hb_0”] <- TRUE
hexamers[names(unlist(sapply(rownames(hexamers), function(X) {agrep(X, Hb, 1, ignore.case=T)}))), “Hb_1”] <- TRUE
hexamers[names(unlist(sapply(rownames(hexamers), function(X) {agrep(X, Hb, 2, ignore.case=T)}))), “Hb_2”] <- TRUE
~~~

~~~
summary(hexamers)
         Hb_0               Hb_1               Hb_2
  Mode :logical     Mode :logical     Mode:logical
FALSE:3154          FALSE:33          TRUE:4096
TRUE :942             TRUE :4063             NA’s:0
NA’s :0                 NA’s :0
~~~

~~~
with(hexamers, rownames(hexamers)[! (Hb_1)])
[1]     “GTTAAA”     “CGACAA”     “GGATAA”     “GTATAA”     “CTACGA”     “TATCGA”     “CGAATA”     “GATATA”
[9]     “CGTATA”     “GTACTA”     “TACCTA”     “ATCGTA”     “CTCGTA”     “TCGTTA”     “TAAAAC”     “TACAAC”
[17]     “ATTTAC”     “AAACCC”     “TAATGC”     “ATCTGC”     “CTAATC”     “ATTCCG”     “CTATCG”     “GATTCG”
[25]     “TACGAT”     “ATCGAT”     “ATCTAT”     “TCGTAT”     “CTAATT”     “TCCATT”     “CCGATT”     “TCGATT”
[33]     “CGATTT”
~~~

## Selection of 40N6 primers

### R code

~~~
acgt <- c(‘A’, ‘C’, ‘G’, ‘T’)
hexamers <- apply(expand.grid(acgt, acgt, acgt, acgt, acgt, acgt), 1, paste, collapse=‘‘)
sample(hexamers,40)
[1]     “CCAGTC”     “CCCTTC”     “TTTTTT”     “CTGTAC”     “TGACCG”     “TGTGAT”     “AACCCT”     “AGGCGG”
[9]     “TCGTCT”     “CTACAA”     “GTACGC”     “CAGAAG”     “GTGTCT”     “GTGTGC”     “AAGACT”     “CGGGTA”
[17]     “AAGAGA”     “GAGGTG”     “GCTCTT”     “GGTGTG”     “GCACGT”     “TGAACT”     “GGGGCG”     “GAGAGG”
[25]     “CCTCAG”     “TAAGTT”     “ATCTGC”     “ACTTAA”     “CACAGC”     “AGATGA”     “GGTAGC”     “AAGGCC”
[33]     “CGCAGG”     “AACCTC”     “CAGTTG”     “ATTCCC”     “AGATGG”     “GCGGAC”     “CTGGCG”     “CTTCAC”
~~~

## Common analysis for all the experiments

### configuration

Use appropriate names instead of xxx (see detailed commands for each experiment)

~~~
library(plyr)
exportInEnv <- function(x) {
    Name <- X
    Value <- get(X)
    .Internal(Sys.setenv(Name, Value))
    cat( paste0(“export “, paste(Name, Value, sep=‘=‘), “\n”))
}
LIBRARY <- ‘xxx’
MOIRAI_USER <- ‘xxx’
MOIRAI_PROJECT <- ‘xxx’
GROUP_SHARED <- ‘xxx’
WORKDIR <- ‘.’
GENE_SYMBOLS <- paste(GROUP_SHARED,
‘annotation/homo_sapiens/gencode-
14/gencode.v14.annotation.genes.bed’, sep=‘/’)
ANNOTATION       <- paste(GROUP_SHARED,
‘annotation/homo_sapiens/100712hg19/100712hg19’, sep=‘/’)
PROCESSED_DATA <- dirname( system( paste( ‘ls -d /osc-fs_home /scratch/moirai/’
                                                                             , MOIRAI_USER, ‘/project/’
                                                                             , MOIRAI_PROJECT, ‘/’
                                                                             , LIBRARY
                                                                             , ‘*/Moirai.config’
                                                                             , sep=‘‘)
                                                                 , intern=TRUE)[1])
l_ply( c(“LIBRARY”, “MOIRAI_USER”, “MOIRAI_PROJECT”,
“GROUP_SHARED”
                         , “WORKDIR”, “GENE_SYMBOLS”, “ANNOTATION”,
“PROCESSED_DATA”)
                 , exportInEnv)
~~~

## Cluster with the PromoterPipeline

### Level 1

Transform the paired-end alignments into level 1 clusters, sort the file and index it. Select only BAM files that contain aligned reads.

~~~
ALIGNED_DATA=$(for BAM in
$PROCESSED_DATA/properly_paired_rmdup/*bam; do samtools
flagstat $BAM | grep -Lq ‘^0 + 0 mapped’ || echo $BAM; done)
level1.py --help | head -n1
level1.py -o /dev/stdout -f 66 -F 516 $ALIGNED_DATA |
   bgzip > $LIBRARY.l1.gz
cat <(zgrep \# -A1 $LIBRARY.l1.gz) <(zgrep -v \#
$LIBRARY.l1.gz | sed ‘1d’ |
    sort --field-separator $’\t’ -k2.4, 2n -k 2.4, 2.4 -k3, 3n
-k4, 4n -k5,5) |
  bgzip |
  sponge $LIBRARY.l1.gz
#tabix -s2 -b3 -e4 $LIBRARY.l1.gz
~~~

### Level 2

Same for level 2 clusters.

Needs a version of level2.py that is more recent than 20120628, where the “Output” message is sent to stderr.

~~~
level2.py --help | head -n1
level2.py -o /dev/stdout -t 0 $LIBRARY.l1.gz |
   bgzip > $LIBRARY.l2.gz
cat <(zgrep \# -A1 $LIBRARY.l2.gz) <(zgrep -v \#
$LIBRARY.l2.gz | sed ‘1d’ |
    sort --field-separator $’t’ -k2.4, 2n -k 2.4, 2.4 -k3, 3n
-k4, 4n -k5,5) |
   bgzip |
   sponge $LIBRARY.l2.gz
#tabix -s2 -b3 -e4 $LIBRARY.l2.gz
~~~

## Intersections

Convert level 1 and 2 files to BED format, and intersect them with pre-defined annotation files.

~~~
function osc2bed {
    zcat $1 |
       grep -v \# |
       sed 1d |
       awk ‘{OFS=“\t”}{print $2, $3, $4, “l1”, “1000”, $5}’
}
function bed2annot {
   bedtools intersect -a $1 -b $ANNOTATION.annot -s -loj |
      awk ‘{OFS=“\t”}{print $1”: “$2” – “$3$6,$10}’ |
      bedtools groupby -g 1 -c 2 -o collapse
}
for LEVEL in l1 l2
do
    osc2bed $LIBRARY.$LEVEL.gz | tee $LIBRARY.$LEVEL.bed |
bed2annot - > $LiBRARY.$LEVEL.annot
done
~~~

## Gene symbols

~~~
function bed2symbols {
   bedtools intersect -a $1 -b $GENE_SYMBOLS -s -loj |
       awk ‘{OFS=“\t”}{print $1”: “$2” – “$3$6, $10}’ |
       bedtools groupby -g 1 -c 2 -o distinct >
$LIBRARY.l2.genes
}
if [$GENE_SYMBOLS]
then
   bed2symbols $LIBRARY.l2.bed > $LIBRARY.l2.genes
fi
~~~

## Analyze of the first experiment: NCms10058

### Configuration

~~~
**library**(plyr)
exportInEnv <- **function**(X) {
   Name <- X
   Value <- get(X)
.Internal(Sys.setenv(Name, Value))
cat( paste0(“export “, paste(Name, Value, sep=‘=‘), “\n”))
}
LIBRARY                  <- ‘NCms10058_1’
MOIRAI_USER         <- ‘nanoCAGE2’
MOIRAI_PROJECT   <- ‘Arnaud’
GROUP_SHARED     <- ‘/osc-fs_home/scratch/gmtu’
WORKDIR                 <- ‘.’
GENE_SYMBOLS     <- paste(GROUP_SHARED, ‘annotation/homo_sapiens/gencode-14/gen code.v14.annotation.genes.bed’, sep=‘/’)
ANNOTATION      <- paste(GROUP_SHARED, ‘annotation/homo_sapiens/100712hg19/100
712hg19’, sep=‘/’)
PROCESSED_DATA <- dirname( system( paste( ‘ls -d /osc-fs_home/scratch/moirai/
                                                                               , MOIRAI_USER
                                                                               , ‘/project/’
                                                                               , MOIRAI_PROJECT
                                                                               , ‘/’
                                                                               , LIBRARY
                                                                               , ‘*/Moirai.config’
                                                                               , sep=‘‘)
                                                                               , intern=TRUE) [1])
l_ply( c(“LIBRARY”, “MOIRAI_USER”, “MOIRAI_PROJECT”, “GROUP_SHARED”
              , “WORKDIR”, “GENE_SYMBOLS”, “ANNOTATION”, “PROCESSED_DATA”)
        , exportInEnv)
~~~

~~~
export LIBRARY=NCms10 058_1
export MOIRAI_USER=nanoCAGE2
export MOIRAI_PROJECT=Arnaud
export GROUP_SHARED=/osc-fs_home/scratch/gmtu
export WORKDIR=.
export GENE_SYMBOLS=/osc-fs_home/scratch/gmtu/annotation/homo_sapiens/gencode
-14/gencode.v14.annotation.genes.bed
export ANNOTATION=/osc-fs_home/scratch/gmtu/annotation/homo_sapiens/100 712hg1
9/100712hg19
export PROCESSED_DATA=/osc-fs_home/scratch/moirai/nanoCAGE2/project/Arnaud/NC
ms10058_1.CAGEscan short-reads.20150625154711
~~~

## Count the reads

~~~
awk ‘/raw/ {print $3}’ $PROCESSED_DATA/text/summary.txt |
    /usr/lib/filo/stats |
    grep ‘Sum’ |
    cut -f2 -d’:’ |
    tr -d ‘[:space:]’ |
    xargs -0 /usr/bin/printf “ # %’d\n”
~~~

~~~
##   # 3608777
~~~

~~~
grep raw $PROCESSED_DATA/text/summary.txt
~~~

~~~
## NCms10058_1.ACAGTG.R1      raw 95519
## NCms10058_1.ACTTGA.R1      raw 76278
## NCms10058_1.ATCACG.R1      raw 53374
## NCms10058_1.CAGATC.R1      raw 103408
## NCms10058_1.CGATGT.R1      raw 73164
## NCms10058_1.CTTGTA.R1      raw 134779
## NCms10058_1.GATCAG.R1      raw 95648
## NCms10058_1.GCCAAT.R1      raw 76012
## NCms10058_1.GGCTAC.R1      raw 56348
## NCms10058_1.TAGCTT.R1      raw 54492
## NCms10058_1.TGACCA.R1      raw 63262
## NCms10058_1.TTAGGC.R1      raw 95230
## NCms10058_1.Undetermined.R1      raw 2631263
~~~

## Analysis with R

### Configuration

~~~
**library** (oscR) # *See https://github.com/charles-plessy/oscR for oscR.*
**if** (compareVersion(sessionInfo()$otherPkgs$oscR$Version,’0.1.1’) < 0) stop(‘o utdated version of oscR.’)
**library** (smallCAGEqc) *# See https://github.com/charles-plessy/smallCAGEqc for smallCAGEqc.*
**if** (compareVersion(sessionInfo()$otherPkgs$smallCAGEqc$Version,’0.6.0’) < 0)
**stop** (‘Outdated version of smallCAGEqc’)
**library** (vegan)
~~~

~~~
## Loading required package: permute
## Loading required package: lattice
## This is vegan 2.0–10
~~~

~~~
**library** (ggplot2)
~~~

### Load data

~~~
l2_NCki <- read.osc(paste(LIBRARY,’l2’,’gz’,sep=‘.’), drop.coord=T, drop.norm
=T)
colnames(l2_NCki) <- sub(‘raw.NCms100 58_1.’,’NCki_’,colnames(l2_NCki))
colSums(l2_NCki)
~~~

~~~
## NCki_HeLa_PS_A NCki_HeLa_PS_B          NCki_HeLa_PS_C NCki_HeLa_RanN6_A NC
ki_HeLa_RanN6_B NCki_HeLa_RanN6_C
##                           11800                           13969                           22764                           14137
                  13556                           10430
##         NCki_THP1_PS_A         NCki_THP1_PS_B         NCki_THP1_PS_C         NCki_THP1_RanN6_A NC
ki_THP1_RanN6_B NCki_THP1_RanN6_C
##                            15157                            15453                            13092                            8708
                     14536                            17122
~~~

### Normalization number of read per sample: l2.sub; libs$genes.sub

In all the 3 libraries used, one contain only few reads tags. The smallest one has 8,708 counts. In order to make meaningful comparisons, all of them are subsapled to 8700 counts.

~~~
l2.sub1 <- t(rrarefy(t(l2_NCki),min(8700)))
colSums(l2.sub1)
~~~

~~~
##     NCki_HeLa_PS_A     NCki_HeLa_PS_B     NCki_HeLa_PS_C     NCki_HeLa_RanN6_A     NC
ki_HeLa_RanN6_B NCki_HeLa_RanN6_C
##                              8700                              8700                              8700                              8700
                      8700                              8700
##       NCki_THP1_PS_A       NCki_THP1_PS_B       NCki_THP1_PS_C NCki_THP1_RanN6_A NC
ki_THP1_RanN6_B NCki_THP1_RanN6_C
##                              8700                              8700                              8700                              8700
                       8700                              8700
~~~

### Moirai statistics

Load the QC data produced by the Moirai workflow with which the libraries were processed. Sort in the same way as the l1 and l2 tables, to allow for easy addition of columns.

~~~
libs <- loadLogs(‘moirai’)
~~~

### Number of clusters

Count the number of unique L2 clusters per libraries after subsampling, and add this to the QC table. Each subsampling will give a different result, but the mean result can be calculated by using the rarefy function at the same scale as the subsampling.

~~~
libs[“l2.sub1”] <- colSums(l2.sub1 > 0)
libs[“l2.sub1.exp”] <- rarefy(t(l2_NCki), min(colSums(l2_NCki)))
~~~

### Richness

Richness should also be calculated on the whole data.

~~~
libs[“r100.l2”] <- rarefy(t(l2_NCki),100)
boxplot(data=libs, r100.l2 ~ group, ylim=c(92,100), las=1)
~~~

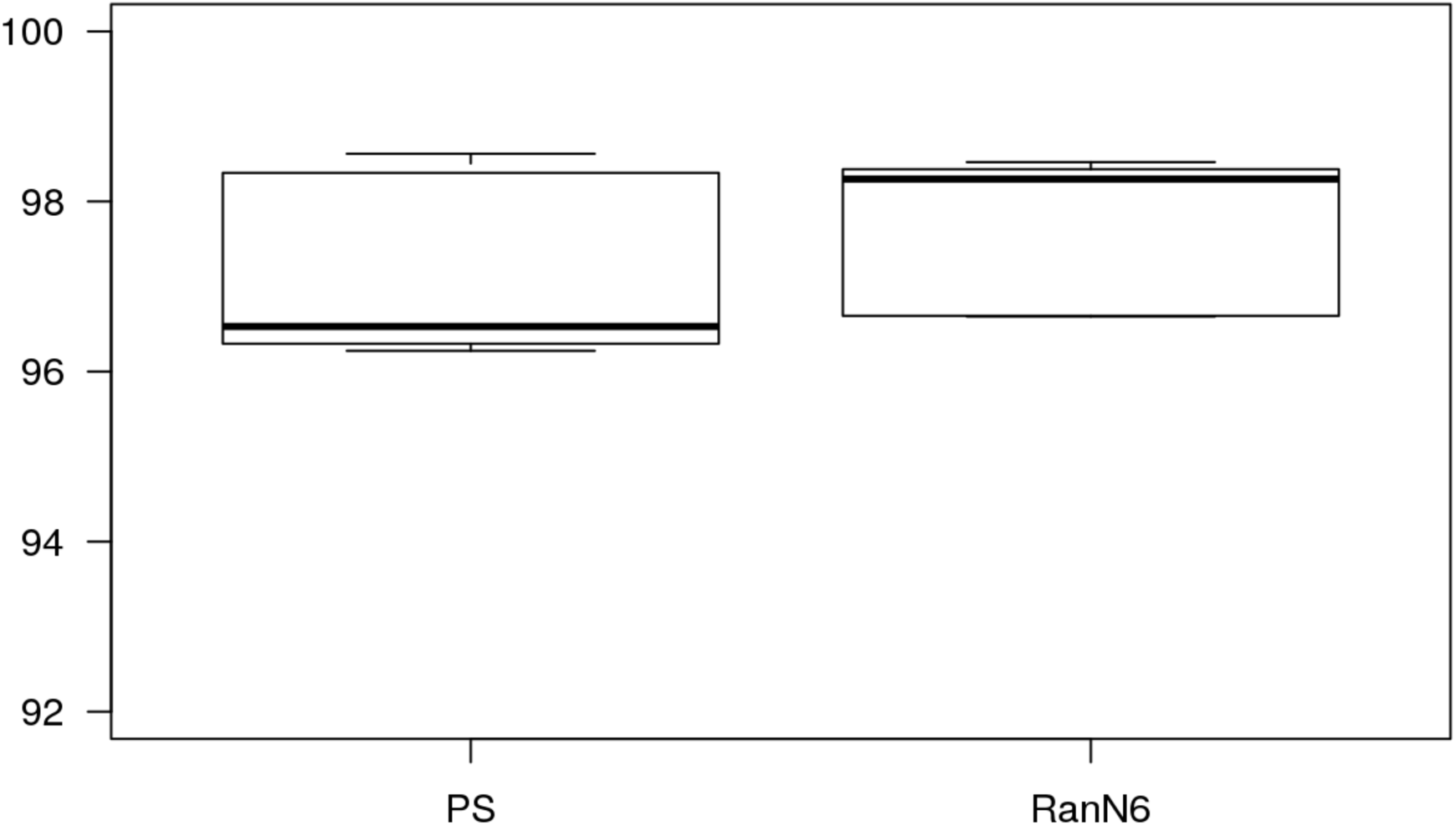

### Hierarchical annotation

Differences of sampling will not bias distort the distribution of reads between annotations, so the non-subsampled library is used here.

~~~
annot.l2 <- read.table(paste(LIBRARY,’l2’,’annot’,sep=‘.’), head=F, col.names
=c(‘id’, ‘feature’), row.names=1)
annot.l2 <- hierarchAnnot(annot.l2)
rownames(libs) <- sub(“HeLa”, “NCki_HeLa”, rownames(libs))
rownames(libs) <- sub(“THP1”, “NCki_THP1”, rownames(libs))
libs <- cbind(libs, t(rowsum(l2_NCki, annot.l2[,’class’])))
libs$samplename <- sub(‘HeLa’, ‘NCki_HeLa’, libs$samplename)
libs$samplename <- sub(‘THP1’, ‘NCki_THP1’, libs$samplename)
~~~

### Gene symbols used normalisation data

~~~
genesymbols <- read.table(paste(LIBRARY,’l2’,’genes’,sep=‘.’), col.names=c(“c
luster”,”symbol”), stringsAsFactors=FALSE)
rownames(genesymbols) <- genesymbols$cluster
g2 <- rowsum(l2_NCki, genesymbols$symbol)
countSymbols <- countSymbols(g2)
libs[colnames(l2_NCki),”genes”] <- (countSymbols)
~~~

Number of genes detected in sub-sample

~~~
l2.sub1 <- data.frame(l2.sub1)
g2.sub1 <- rowsum(l2.sub1, genesymbols$symbol)
countSymbols. sub1 <- countSymbols(g2.sub1)
libs[colnames(l2.sub1), “genes.sub1”] <- (countSymbols.sub1)
~~~

### Table record

save the different tables produced for later analysis

~~~
write.table(l2_NCki, “l2_NCki_1.txt”, sep = “\t”, quote=FALSE)
write.table(l2.sub1, “l2.sub1_NCki_1.txt”, sep = “\t”, quote=FALSE)
write.table(g2.sub1, ‘g2.sub1_NCki_1.txt’, sep=“\t”, quote=F)
write.table(libs, ‘libs_NCki_1.txt’, sep=“\t”, quote=F)
~~~

## Analyze of the second experiment: NC12

### configuration

~~~
**library** (plyr)
exportInEnv <- **function**(X) {
   Name <- X
   Value <- get(X)
   .Internal(Sys.setenv(Name, Value))
   cat( paste0 (“export”, paste(Name, Value, sep=‘=‘), “\n”))
^}^
LIBRARY                   <- ‘NC12_1’
MOIRAI_USER          <- ‘nanoCAGE2’
MOIRAI_PROJECT   <- ‘Arnaud’
GROUP_SHARED     <- ‘/osc-fs_home/scratch/gmtu’
WORKDIR                  <- ‘.’
GENE_SYMBOLS      <- paste(GROUP_SHARED, ‘annotation/homo_sapiens/gencode-14/gen
code.v14.annotation.genes.bed’, sep=‘/’)
ANNOTATION         <- paste(GROUP_SHARED, ‘annotation/homo_sapiens/100 712hg19/100
712hg19’, sep=‘/’)
PROCESSED_DATA <- dirname( system( paste( ‘ls -d /osc-fs_home/scratch/moirai/
                                                                               , MOIRAI_USER
                                                                               , ‘/project/’
                                                                               , MOIRAI_PROJECT
                                                                               , ‘/’
                                                                               , LIBRARY
                                                                               , ‘*/Moirai.config’
                                                                               , sep=‘‘)
                                                                     , intern=TRUE)[1])
l_ply( c(“LIBRARY”, “MOIRAI_USER”, “MOIRAI_PROJECT”, “GROUP_SHARED”
             , “WORKDIR”, “GENE_SYMBOLS”, “ANNOTATION”, “PROCESSED_DATA”)
       , exportInEnv)
~~~

~~~
export LIBRARY=NC12_1
export MOIRAI_USER=nanoCAGE2
export MOIRAI_PROJECT=Arnaud
export GROUP_SHARED=/osc-fs_home/scratch/gmtu
export WORKDIR=.
export GENE_SYMBOLS=/osc-fs_home/scratch/gmtu/annotation/homo_sapiens/gencode
-14/gencode.v14.annotation.genes.bed
export ANNOTATION=/osc-fs_home/scratch/gmtu/annotation/homo_sapiens/100 712hg1 9/100712hg19
export PROCESSED_DATA=/osc-fs_home/scratch/moirai/nanoCAGE2/project/Arnaud/NC
12_1.CAGEscan_short-reads.20150629125015
~~~

## Count the reads

~~~
awk ‘/raw/ {print $3}’ $PROCESSED_DATA/text/summary.txt |
     /usr/lib/filo/stats |
     grep ‘Sum’ |
     cut -f2 -d’:’ |
     tr -d ‘[:space:]’ |
     xargs -0 /usr/bin/printf “# %’d\n”
~~~

~~~
##     # 3450701
~~~

~~~
grep raw $PROCESSED_DATA/text/summary.txt
~~~

~~~
## NC12_1.ACAGTG.R1 raw 340256
## NC12_1.ATCACG.R1 raw 437139
## NC12_1.CGATGT.R1 raw 274252
## NC12_1.GCCAAT.R1 raw 390496
## NC12_1.TGACCA.R1 raw 287340
## NC12_1.TTAGGC.R1 raw 316502
## NC12_1.Undetermined.R1         raw 1404716
~~~

## Analysis with R

### Configuration

~~~
**library** (oscR) *# See https://github.com/charles-plessy/oscR for oscR.*
**if** (compareVersion(sessionInfo()$otherPkgs$oscR$Version,’0.1.1’) < 0) **stop**(‘O
utdated version of oscR.’)
**library** (smallCAGEqc) # *See https://github.com/charles-plessy/smallCAGEqc for smallCAGEqc.*
**if** (compareVersion(sessionInfo()$otherPkgs$smallCAGEqc$Version,’0.6.0’) < 0)
**stop**(‘Outdated version of smallCAGEqc’)
**library** (vegan)
~~~

~~~
**library** (ggplot2)
~~~

### Load data

~~~
l2_NC12 <- read.osc(paste(LIBRARY, ‘l2’, ‘gz’, sep=‘.’), drop.coord=T, drop.norm
=T)
colnames(l2_NC12) <- sub(‘raw.NC12_1’, ‘NC12’, colnames(l2_NC12))
colSums(l2_NC12)
~~~

~~~
##    NC12.HeLa_4 0N6_A    NC12.HeLa_40N6_B    NC12.HeLa_40N6_C    NC12.HeLa_PS_A
   NC12.HeLa_PS_B    NC12.HeLa_PS_C
##                             12154                             17411                             20790                             24065
27215                             54835
## NC12.HeLa_RanN6_A NC12.HeLa_RanN6_B NC12.HeLa_RanN6_C    NC12.THP1_40N6_A     N
C12.THP1_40N6_B NC12.THP1_40N6_C
##                            10944                            35582                            23215                            9271                            15299                            15775
##      NC12.THP1_PS_A      NC12.THP1_PS_B      NC12.THP1_PS_C NC12.THP1_RanN6_A NC
12.THP1_RanN6_B NC12.THP1_RanN6_C
##                            21303                            23454                            37395                            13356
                 58890                            34922
~~~

### Normalization number of read per sample: l2.sub; libs$genes.sub

~~~
l2.sub1 <- t(rrarefy(t(l2_NC12),min(8700)))
colSums(l2.sub1)
~~~

~~~
##   NC12.HeLa_4 0N6_A    NC12.HeLa_40N6_B   NC12.HeLa_40N6_C     NC12.HeLa_PS_A
NC12.HeLa_PS_B      NC12.HeLa_PS_C
##                           8700                           8700                           8700                           8700
                      8700                           8700
## NC12.HeLa_RanN6_A NC12.HeLa_RanN6_B NC12.HeLa_RanN6_C    NC12.THP1_40N6_A    N
C12.THP1_40N6_B    NC12.THP1_40N6_C
##                              8700                              8700                              8700                              8700
                        8700                              8700
##       NC12.THP1_PS_A       NC12.THP1_PS_B       NC12.THP1_PS_C NC12.THP1_RanN6_A NC       
12.THP1_RanN6_B NC12.THP1_RanN6_C
##                              8700                              8700                              8700                              8700
                       8700                              8700
~~~

### Moirai statistics

~~~
libs <- loadLogs(‘moirai’)
rownames (libs) <- sub(‘HeLa’, ‘NC12.HeLa’, rownames(libs))
rownames (libs) <- sub(‘THP1’, ‘NC12.THP1’, rownames(libs))
~~~

### Number of clusters

~~~
libs[“l2.sub1”]         <- colSums(l2.sub1 > 0)
libs[“l2.sub1.exp”]  <- rarefy(t(l2_NC12), min(colSums(l2_NC12)))
~~~

### Richness

Richness should also be calculated on the whole data.

~~~
libs[“r100.l2”] <- rarefy (t(12_NC12), 100)
boxplot (data=libs, r100.12 ~ group, ylim=c (92, 100), las=1)
~~~

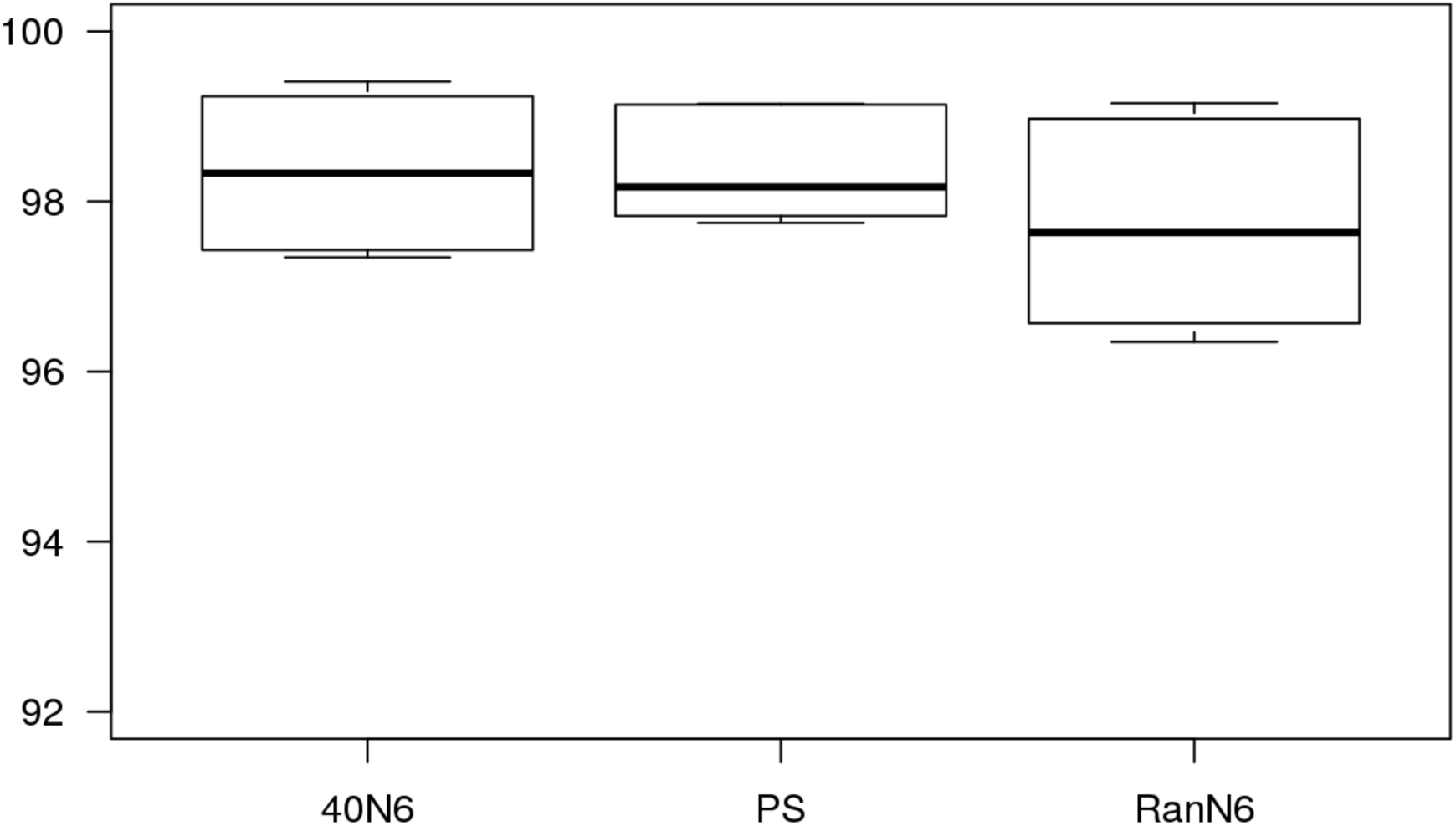

### Hierarchical annotation

~~~
annot.l2 <- read.table(paste(LIBRARY,’l2’,’annot’,sep=‘.’), head=F, col.names
=c(‘id’, ‘feature’), row.names=1)
annot.l2 <- hierarchAnnot(annot.l2)
libs <- cbind(libs, t(rowsum(l2_NC12, annot.l2[,’class’])))
libs$samplename <- sub(‘HeLa’, ‘NC12_HeLa’, libs$samplename)
libs$samplename <- sub(‘THP1’, ‘NC12_THP1’, libs$samplename)
~~~

### Gene symbols used normalisation data

~~~
genesymbols <- read.table(paste(LIBRARY,’l2’,’genes’,sep=‘.’), col.names=c(“c
luster”,”symbol”), stringsAsFactors=FALSE)
rownames(genesymbols) <- genesymbols$cluster
g2 <- rowsum(l2_NC12, genesymbols$symbol)
countSymbols <- countSymbols(g2)
libs[colnames(l2_NC12),”genes”] <- (countSymbols)
~~~

Number of genes detected in sub-sample

~~~
l2.sub1 <- data.frame(l2.sub1)
g2.sub1 <- rowsum(l2.sub1, genesymbols$symbol)
countSymbols.sub1 <- countSymbols(g2.sub1)
libs[colnames(l2.sub1), “genes.sub1”] <- (countSymbols.sub1)
~~~

### Table record

save the different tables produced for later analysis

~~~
write.table(l2_NC12, “l2_NC12_1.txt”, sep = “\t”, quote=FALSE)
write.table(l2.sub1, “l2.sub1_NC12_1.txt”, sep = “\t”, quote=FALSE)
write.table(g2.sub1, ‘g2.sub1_NC12_1.txt’, sep=“\t”, quote=F)
write.table(libs, ‘libs_NC12_1.txt’, sep=“\t”, quote=F)
~~~

## Analyze of the third experiment: NC17

### Configuration

~~~
**library** (plyr)
exportInEnv <- **function**(X) {
   Name <- X
   Value <- get(X)
   .Internal(Sys.setenv(Name, Value))
cat( paste0(“export “, paste(Name, Value, sep=‘=‘), “\n”))
^}^
LIBRARY                   <- ‘NC16–17_1’
MOIRAI_USER          <- ‘nanoCAGE2’
MOIRAI_PROJECT   <- ‘Arnaud’
GROUP_SHARED     <- ‘/osc-fs_home/scratch/gmtu’
WORKDIR                  <- ‘.’
GENE_SYMBOLS      <- paste(GROUP_SHARED, ‘annotation/homo_sapiens/gencode-14/gen
code.v14.annotation.genes.bed’, sep=‘/’)
ANNOTATION        <- paste(GROUP_SHARED, ‘annotation/homo_sapiens/100712hg19/100
712hg19’, sep=‘/’)
PROCESSED_DATA <- dirname( system( paste( ‘ls -d /osc-fs_home/scratch/moirai/
                                                                               , MOIRAI_USER ‘/project/’
                                                                               , MOIRAI_PROJECT ‘/’
                                                                               , LIBRARY
                                                                               , ‘*/Moirai.config’ sep=‘‘)
                                                                               , intern=TRUE) [1])
l_ply( c(“LIBRARY”, “MOIRAI_USER”, “MOIRAI_PROJECT”, “GROUP_SHARED”
              , “WORKDIR”, “GENE_SYMBOLS”, “ANNOTATION”, “PROCESSED_DATA”)
          , exportInEnv)
~~~

~~~
export LIBRARY=NC16–17_1
export MOIRAI_USER=nanoCAGE2
export MOIRAI_PROJECT=Arnaud
export GROUP_SHARED=/osc-fs_home/scratch/gmtu
export WORKDIR=.
export GENE_SYMBOLS=/osc-fs_home/scratch/gmtu/annotation/homo_sapiens/gencode
-14/gencode.v14.annotation.genes.bed
export ANNOTATION=/osc-fs_home/scratch/gmtu/annotation/homo_sapiens/100 712hg1
9/100712hg19
export PROCESSED_DATA=/osc-fs_home/scratch/moirai/nanoCAGE2/project/Arnaud/NC
16–17_1.CAGEscan_short-reads.20150625154740
~~~

### Count the reads

~~~
awk ‘/raw/ {print $3}’ $PROCESSED_DATA/text/summary.txt |
/usr/lib/filo/stats |
grep ‘Sum’ |
cut -f2 -d’:’ |
tr -d ‘[:space:]’ |
xargs -0 /usr/bin/printf “ # %’d\n”
~~~

~~~
##   # 4821156
~~~

~~~
grep raw $PROCESSED_DATA/text/summary.txt
~~~

~~~
## NC16–17_1.ACAGTG.R1 raw 211404
## NC16–17_1.ACTTGA.R1 raw 189074
## NC16–17_1.ATCACG.R1 raw 544817
## NC16–17_1.CAGATC.R1 raw 214188
## NC16–17_1.CGATGT.R1 raw 490410
## NC16–17_1.CTTGTA.R1 raw 308921
## NC16–17_1.GATCAG.R1 raw 167839
## NC16–17_1.GCCAAT.R1 raw 620406
## NC16–17_1.GGCTAC.R1 raw 150422
## NC16–17_1.TAGCTT.R1 raw 200755
## NC16–17_1.TGACCA.R1 raw 386420
## NC16–17_1.TTAGGC.R1 raw 368814
## NC16–17_1.Undetermined.R1         raw 967686
~~~

### Analysis with R

~~~
**library**(oscR) # *See https://github.com/charles-plessy/oscR for oscR.*
**if** (compareVersion(sessionInfo()$otherPkgs$oscR$Version,’0.1.1’) < 0) **stop**(‘O
utdated version of oscR.’)
**library**(smallCAGEqc) *# See https://github.com/charles-plessy/smallCAGEqc for smallCAGEqc.*
**if** (compareVersion(sessionInfo()$otherPkgs$smallCAGEqc$Version,’0.6.0’) < 0)
**stop** (‘Outdated version of smallCAGEqc’)
**library** (vegan)
~~~

~~~
**library**(ggplot2)
**library**(pvclust)
~~~

### Load data

~~~
l2_NC17 <- read.osc(paste(LIBRARY,’l2’,’gz’,sep=‘.’), drop.coord=T, drop.norm =T)
colnames(l2_NC17) <- sub(‘raw.NC16.17_1.17’, ‘NC17’, colnames(l2_NC17))
colSums(l2_NC17)
~~~

~~~
##    NC17_HeLa_10PS_A    NC17_HeLa_10PS_B    NC17_HeLa_10PS_C NC17_HeLa_20PS1_A NC
17_HeLa_20PS1_B NC17_HeLa_20PS1_C
##                           31006                           29327                           34781                           29549
                   18858                           18469
##   NC17_HeLa_20PS2_A   NC17_HeLa_20PS2_B   NC17_HeLa_20PS2_C NC
17_HeLa_20PS3_A NC 17_HeLa_20PS3_B NC17_HeLa_20PS3_C
##                    23579                    15882                    18592                    26289
              15389                    15712
##       NC17_HeLa_PS_A       NC17_HeLa_PS_B       NC17_HeLa_PS_C NC17_HeLa_RanN6_A NC
17_HeLa_RanN6_B NC17_HeLa_RanN6_C
##                            29038                            21308                            29123                            44255
                   17650                            21824
##   NC17_THP1_10PS_A   NC17_THP1_10PS_B   NC17_THP1_10PS_C NC17_THP1_20PS1_A NC
17_THP1_20PS1_B NC17_THP1_20PS1_C
##                              26158                              19394                              28814                              17733
                  14452                              19870
##    NC17_THP1_20PS2_A    NC17_THP1_20PS2_B    NC17_THP1_20PS2_C NC17_THP1_20PS3_A NC
17_THP1_20PS3_B NC17_THP1_20PS3_C
##                        19562                        11486                        21205                        23229
                 21447                        17429
##     NC17_THP1_PS_A     NC17_THP1_PS_B     NC17_THP1_PS_C NC17_THP1_RanN6_A NC
17_THP1_RanN6_B     NC17_THP1_RanN6_C
##                        24370                        18173                        20788                        20236
                   14661                        22048
~~~

### Normalization number of read per sample: l2.sub; libs$genes.sub

~~~
l2.sub1 <- t(rrarefy(t(l2_NC17),min(8700)))
colSums(l2.sub1)
~~~

~~~
##   NC17_HeLa_10PS_A   NC17_HeLa_10PS_B   NC17_HeLa_10PS_C NC17_HeLa_20PS1_A NC
17_HeLa_20PS1_B NC17_HeLa_20PS1_C
##                           8700                           8700                           8700                           8700
                    8700                           8700
## NC17_HeLa_20PS2_A NC17_HeLa_20PS2_B NC17_HeLa_20PS2_C NC17_HeLa_20PS3_A NC
17_HeLa_20PS3_B NC17_HeLa_20PS3_C
##                           8700                           8700                           8700                           8700
                    8700                           8700
##     NC17_HeLa_PS_A     NC17_HeLa_PS_B     NC17_HeLa_PS_C NC17_HeLa_RanN6_A NC
17_HeLa_RanN6_B NC17_HeLa_RanN6_C
##                    8700                    8700                    8700                    8700
               8700                    8700
##   NC17_THP1_10PS_A   NC17_THP1_10PS_B NC17_THP1_10PS_C NC17_THP1_20PS1_A NC
17_THP1_20PS1_B NC17_THP1_20PS1_C
##                     8700                     8700                     8700                     8700
                  8700                     8700
## NC17_THP1_20PS2_A NC17_THP1_20PS2_B NC17_THP1_20PS2_C NC17_THP1_20PS3_A NC
17_THP1_20PS3_B NC17_THP1_20PS3_C
##                     8700                     8700                     8700                     8700
                8700                     8700
##     NC17_THP1_PS_A     NC17_THP1_PS_B     NC17_THP1_PS_C NC17_THP1_RanN6_A NC
17_THP1_RanN6_B NC17_THP1_RanN6_C
##                    8700                    8700                    8700                    8700
               8700                    8700
~~~

### Moirai statistics

~~~
libs <- loadLogs(‘moirai’)
~~~

### Number of clusters

~~~
libs[“l2.sub1”]         <- colSums(l2.sub1 > 0)
libs[“l2.sub1.exp”]  <- rarefy(t(l2_NC17), min(colSums(l2_NC17)))
~~~

### Richness

Richness should also be calculated on the whole data.

~~~
libs[“r100.l2”] <- rarefy(t(l2_NC17),100)
~~~

~~~
boxplot(data=libs, r100.l2 ~ group, ylim=c(92,100), las=1)
~~~

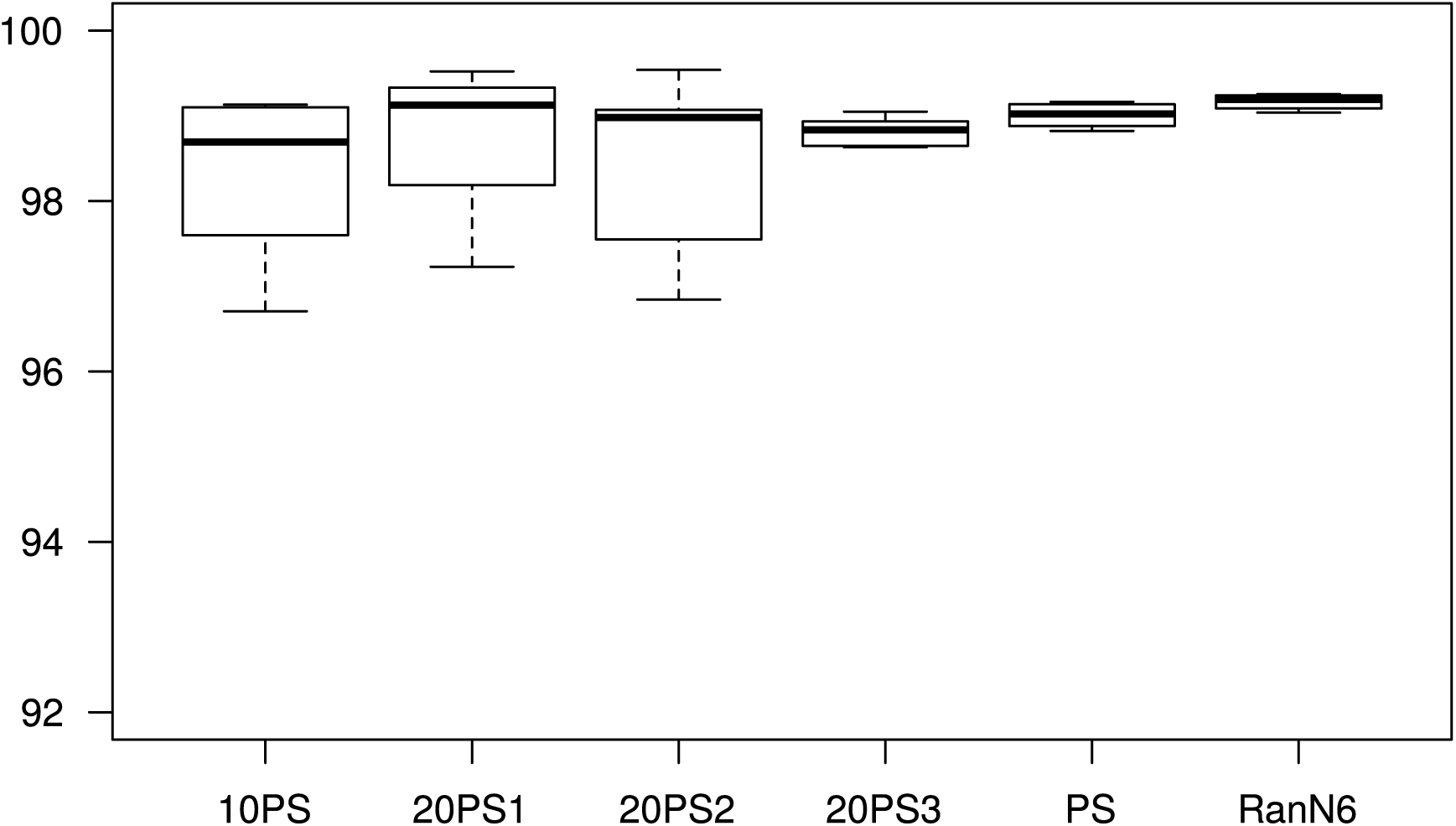

### Hierarchical annotation

~~~
annot.l2 <- read.table(paste(LIBRARY,’l2’,’annot’,sep=‘.’), head=F, col.names
=c(‘id’, ‘feature’), row.names=1)
annot.l2 <- hierarchAnnot(annot.l2)
rownames(libs) <- sub(“17_”, “NC17_”, rownames(libs))
libs <- cbind(libs, t(rowsum(l2_NC17, annot.l2[,’class’])))
libs$samplename <- sub(‘17_’, ‘NC17_’, libs$samplename)
~~~

### Gene symbols used normalisation data

~~~
genesymbols <- read.table(paste(LIBRARY,’l2’,’genes’,sep=‘.’), col.names=c(“c
luster”,”symbol”), stringsAsFactors=FALSE)
rownames(genesymbols) <- genesymbols$cluster
g2 <- rowsum(l2_NC17, genesymbols$symbol)
countSymbols <- countSymbols(g2)
libs[colnames(l2_NC17),”genes”] <- (countSymbols)
~~~

Number of genes detected in sub-sample

~~~
l2.sub1 <- data.frame(l2.sub1)
g2.sub1 <- rowsum(l2.sub1, genesymbols$symbol)
countSymbols.sub1 <- countSymbols(g2.sub1)
libs[colnames(l2.sub1),”genes.sub1”] <- (countSymbols.sub1)
~~~

## Comparison trancriptome

~~~
m2 <- data.frame(
    HeLa_RanN6 = rowMeans (g2[, c(‘NC17_HeLa_RanN6_A’,        ‘NC17_HeLa_RanN6_B’,
    ‘NC17_HeLa_RanN6_C’)]),
    HeLa_PS = rowMeans(g2[, c(‘NC17_HeLa_PS_A’, ‘NC17_HeLa_PS_B’, ‘NC17_HeLa_PS
_C’)]),
    HeLa_20PS3 = rowMeans(g2[, c(‘NC17_HeLa_20PS3_A’, ‘NC17_HeLa_20PS3_B’, ‘NC1
7_HeLa_20PS3_C’)]),
HeLa_20PS1 = rowMeans(g2[, c(‘NC17_HeLa_20PS1_A’, ‘NC17_HeLa_20PS1_B’, ‘NC1
7_HeLa_20PS1_C’)]),
HeLa_20PS2 = rowMeans(g2[, c(‘NC17_HeLa_20PS2_A’, ‘NC17_HeLa_20PS2_B’, ‘NC1
7_HeLa_20PS2_C’)]),
    HeLa_10PS = rowMeans(g2[, c(‘NC17_HeLa_10PS_A’, ‘NC17_HeLa_10PS_B’, ‘NC17_H
eLa_10PS_C’)]),
    THP1_RanN6 = rowMeans(g2[, c(‘NC17_THP1_RanN6_A’,     ‘NC17_THP1_RanN6_B’, ‘NC17_THP1_RanN6_C’)]),
    THP1_PS = rowMeans(g2[, c(‘NC17_THP1_PS_A’, ‘NC17_THP1_PS_B’, ‘NC17_THP1_PS
_C’)]),
    THP1_20PS3 = rowMeans(g2[, c(‘NC17_THP1_20PS3_A’, ‘NC17_THP1_20PS3_B’, ‘NC1
7_THP1_20PS3_C’)]),
    THP1_20PS1 = rowMeans(g2[, c(‘NC17_THP1_20PS1_A’, ‘NC17_THP1_20PS1_B’, ‘NC1
7_THP1_20PS1_C’)]),
    THP1_20PS2 = rowMeans(g2[, c(‘NC17_THP1_20PS2_A’, ‘NC17_THP1_20PS2_B’, ‘NC1
7_THP1_20PS2_C’)]),
THP1_10PS = rowMeans(g2[, c(‘NC17_THP1_10PS_A’, ‘NC17_THP1_10PS_B’, ‘NC17_T
HP1_10PS_C’)])
)
~~~

~~~
results <- pvclust(m2)
~~~

~~~
## Bootstrap (r = 0.5)… Done.
## Bootstrap (r = 0.6)… Done.
## Bootstrap (r = 0.7)… Done.
## Bootstrap (r = 0.8)… Done.
## Bootstrap (r = 0.9)… Done.
## Bootstrap (r = 1.0)… Done.
## Bootstrap (r = 1.1)… Done.
## Bootstrap (r = 1.2)… Done.
## Bootstrap (r = 1.3)… Done.
## Bootstrap (r = 1.4)… Done.
~~~

~~~
plot(results)
pvrect(results, alpha=0.95)
~~~

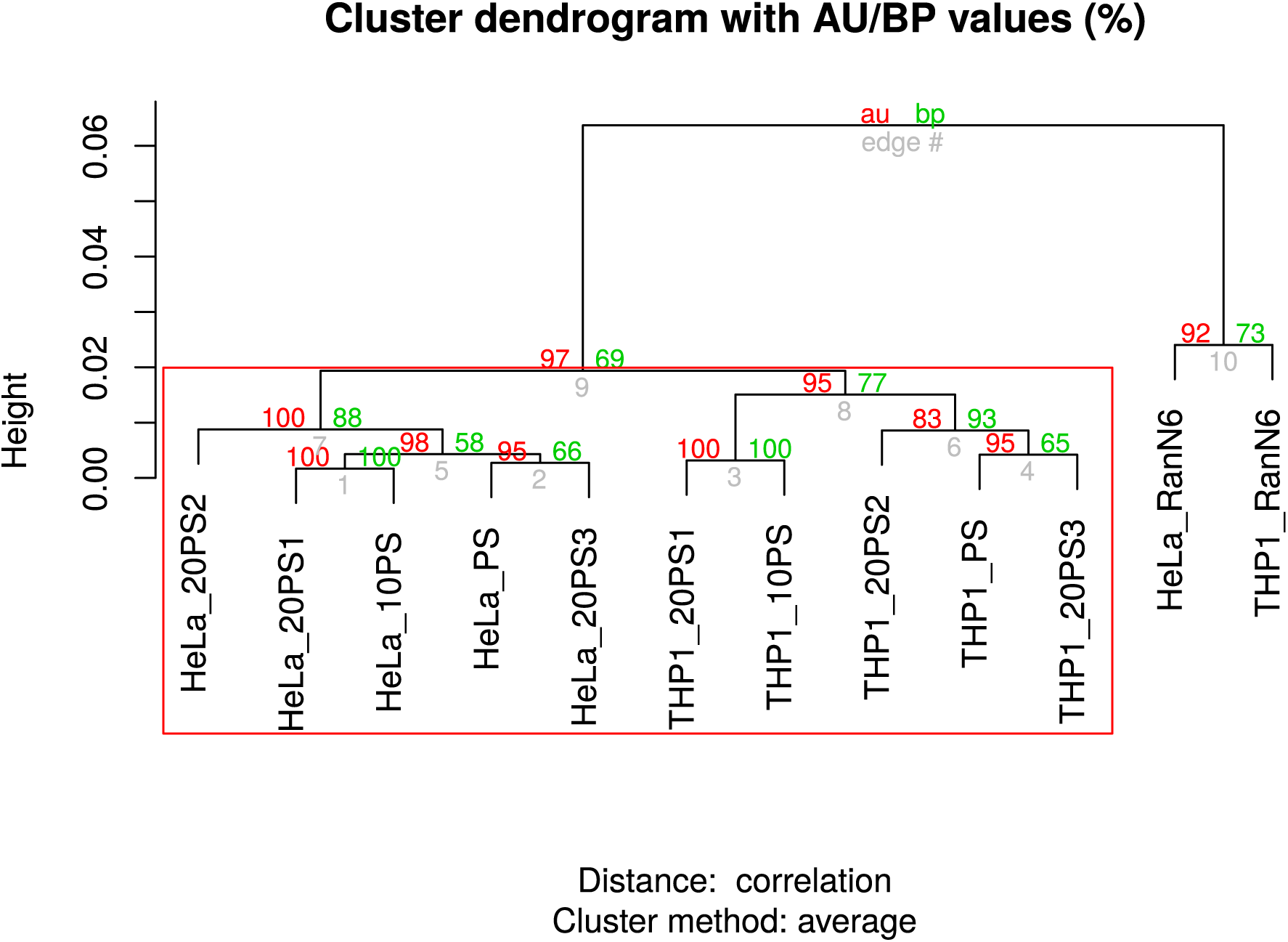

## Table record

save the different tables produced for later analysis

~~~
write.table(l2_NC17, “l2_NC17_1.txt”, sep = “\t”, quote=FALSE)
write.table(l2.sub1, “l2.sub1_NC17_1.txt”, sep = “\t”, quote=FALSE)
write.table(g2.sub1, ‘g2.sub1_NC17_1.txt’, sep=“\t”, quote=F)
write.table(libs, ‘libs_NC17_1.txt’, sep=“\t”, quote=F)
write.table(m2, “m2_NC17_1.txt”, sep = “\t”, quote = FALSE)
~~~

## Analyze of the 3 experiments

### Analysis with R

#### Configuration

~~~
**library**(oscR) # *See https://github.com/charles-plessy/oscR*
**if** (compareVersion(sessionInfo()$otherPkgs$oscR$Version,’0.1.1’) < 0)
   **stop**(‘Out of date oscR library’)
**library**(smallCAGEqc) *# See https://github.com/charles-plessy/smallCAGEqc*
**if** (compareVersion(sessionInfo()$otherPkgs$smallCAGEqc$Version,’0.6.0’) < 0)
   **stop**(‘Out of date smallCAGEqc library’)
**library**(gdata)
~~~

~~~
## gdata: read.xls support for ‘XLS’ (Excel 97–2004) files ENABLED.
##
## gdata: read.xls support for ‘XLSX’ (Excel 2007+) files ENABLED.
##
## Attaching package: ‘gdata’
##
## The following object is masked from ‘package:stats’:
##
##        nobs
##
## The following object is masked from ‘package:utils’:
##
##       object.size
~~~

~~~
**library**(vegan)
~~~

~~~
**library**(ggplot2)
~~~

#### Load the data

~~~
libs_NC12 <- read.table(“libs_NC12_1.txt”, sep=“\t”, head=T)
libs_NCki <- read.table(“libs_NCki_1.txt”, sep=“\t”, head=T)
libs_NC17 <- read.table(“libs_NC17_1.txt”, sep=“\t”, head=T)
~~~

#### Merge 3 tables

The data coming from the 3 experiments are merged in one table to analyzed them together

~~~
rownames(libs_NC12) <- sub(‘NC12.’, ‘NC12_’, rownames(libs_NC12))
~~~

~~~
libs <- rbind(libs_NC12, libs_NC17, libs_NCki)
~~~

##### Add the celltype

~~~
libs$celltype <- libs$samplename
libs$celltype <- sub(‘NC.._’, libs$celltype)
libs$celltype <- sub(‘_.*’, libs$celltype)
libs$celltype <- factor(libs$celltype)
~~~

#### Figure S2

Modification of the table libs (group by triplicates)

~~~
libs2 <- libs
libs2$group <-libs2$samplename
libs2$group <- sub(‘_.$’, ‘‘, libs2$group)
libs2$group <- factor(libs2$group)
~~~

~~~
plotAnnot(libs2, ‘all’, ‘pseudo-random primers’) + theme_bw()
~~~

~~~
## Using group as id variables
## Using group as id variables
~~~

~~~
## Warning: Stacking not well defined when ymin != 0
~~~

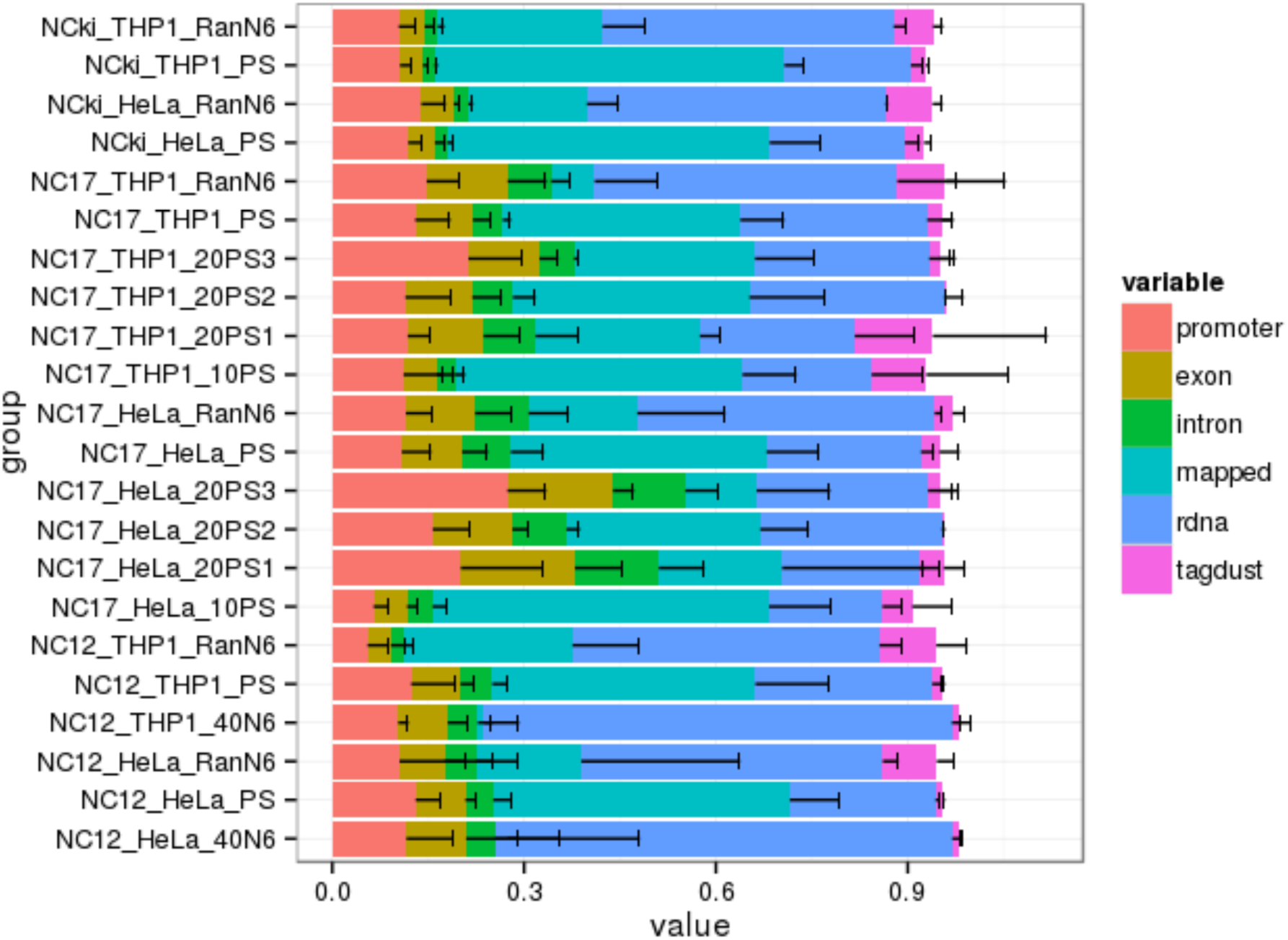

~~~
libs <- libs[grep(‘_RanN6|_PS|_40N6’, libs$samplename, value=T),]
libs <- drop.levels(libs)
write.table(libs, “libs.txt”, sep=“\t”, quote=F)
~~~

#### Impact rDNA and artefacts

Calculate means by triplicate

~~~
libm <- with (libs
, data.frame(samplename, group, celltype
, promoter = promoter / extracted
, exon = exon / extracted
, intron = intron/extracted
, unknown = unknown / extracted
, rDNA = rdna / extracted
, artefacts = tagdust / extracted
))
libm$triplicates <- sub(‘_.$’, ‘‘, libm$samplename)
libm <- aggregate(libm[,c(‘rDNA’,’artefacts’)], list(libm$triplicates), mean)
libm$artefact1000 <- (libm$artefacts)*1000
libm$rDNA1000 <- (libm$rDNA)*1000
libm$group <- libm$Group.1
libm$group <- sub(‘NC.._’, “,libm$group)
~~~

Draw graph

~~~
dotsize <- mean(libm$artefact1000) /5
p <- ggplot(libm, aes(y=artefact1000, x=group)) +
stat_summary(fun.y=mean, fun.ymin=mean, fun.ymax=mean,
geom=“crossbar”, color=“gray”) +
              geom_dotplot(aes(fill=group), binaxis=‘y’, binwidth=1,
dotsize=dotsize, stackdir=‘center’) +
             theme_bw() +
     theme(axis.text.x = element_text(size=13, angle=90)) +
     theme(axis.text.y = element_text(size=13)) +
     theme(axis.title.x = element_blank())+
     theme(axis.title.y = element_text(size=14))+
     scale_y_continuous(breaks =c(0, 50, 100, 150, 200), limits= c(0,200), lab
     els=c(“0”, “0.05”, “0.1”, “0.15”, “0.2”)) +
              ylab(“aretfacts / extracted”)
p + theme(legend.position=“none”)
~~~

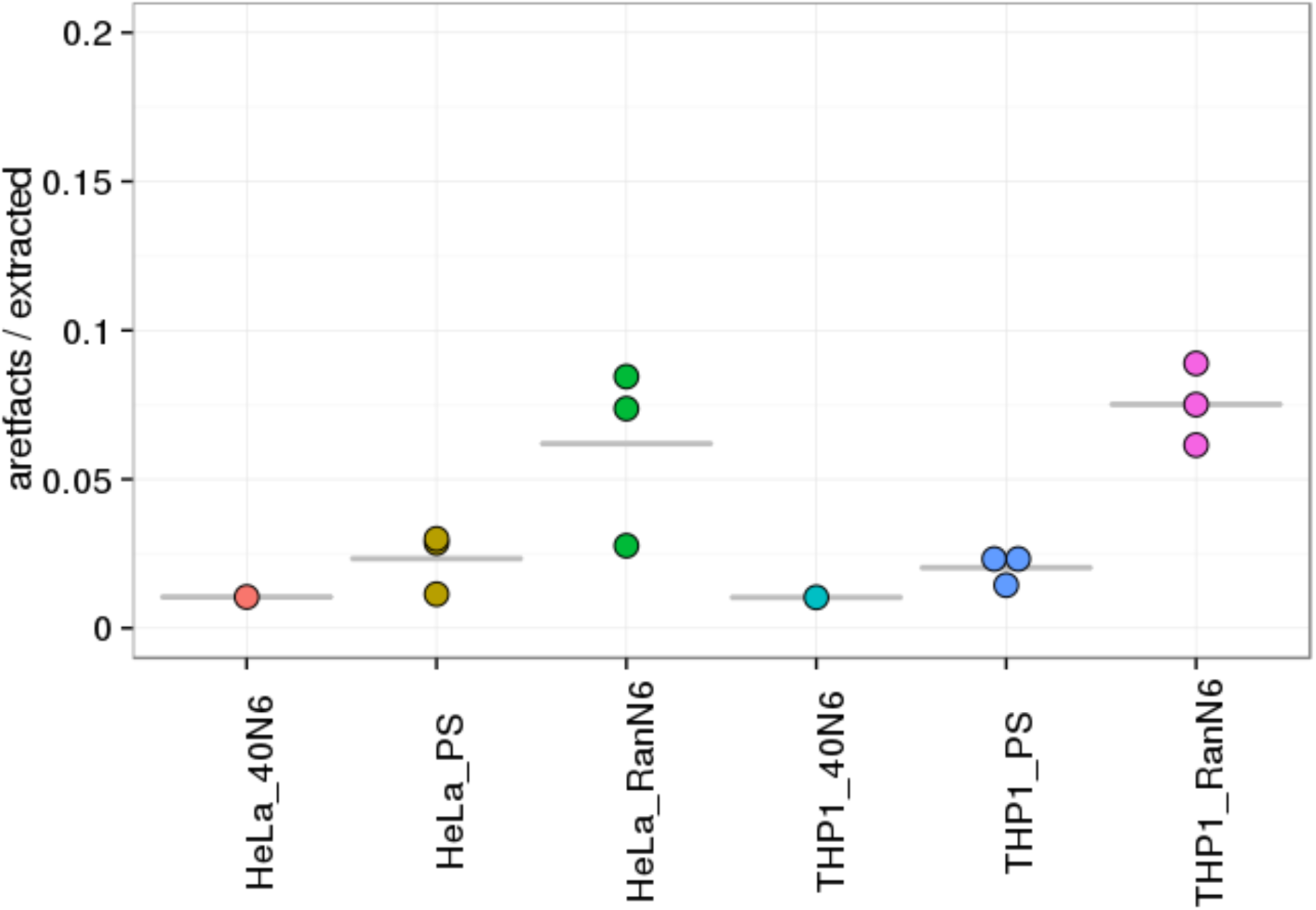

~~~
dotsize <- mean(libm$rDNA1000)/10
P <- ggplot(libm, aes(y=rDNA100 0, x=group)) +
stat_summary(fun.y=mean, fun.ymin=mean, fun.ymax=mean,
geom=“crossbar”, color=“gray”) +
              geom_dotplot(aes(fill=group), binaxis=‘y’, binwidth=1,
dotsize=dotsize, stackdir=‘center’) +
             theme_bw() +
     theme(axis.text.x = element_text(size=13, angle=90)) +
     theme(axis.text.y = element_text(size=13)) +
     theme(axis.title.x = element_blank())+
     theme(axis.title.y = element_text(size=14))+
     scale_y_continuous(limits=c(0,900), breaks =c(0, 200, 400, 600, 800), lab
els=c(“0”, “0.2”, “0.4”, “0.6”, “0.8”)) +
         ylab(“rDNA / extracted”)
p + theme(legend.position=“none”)
~~~

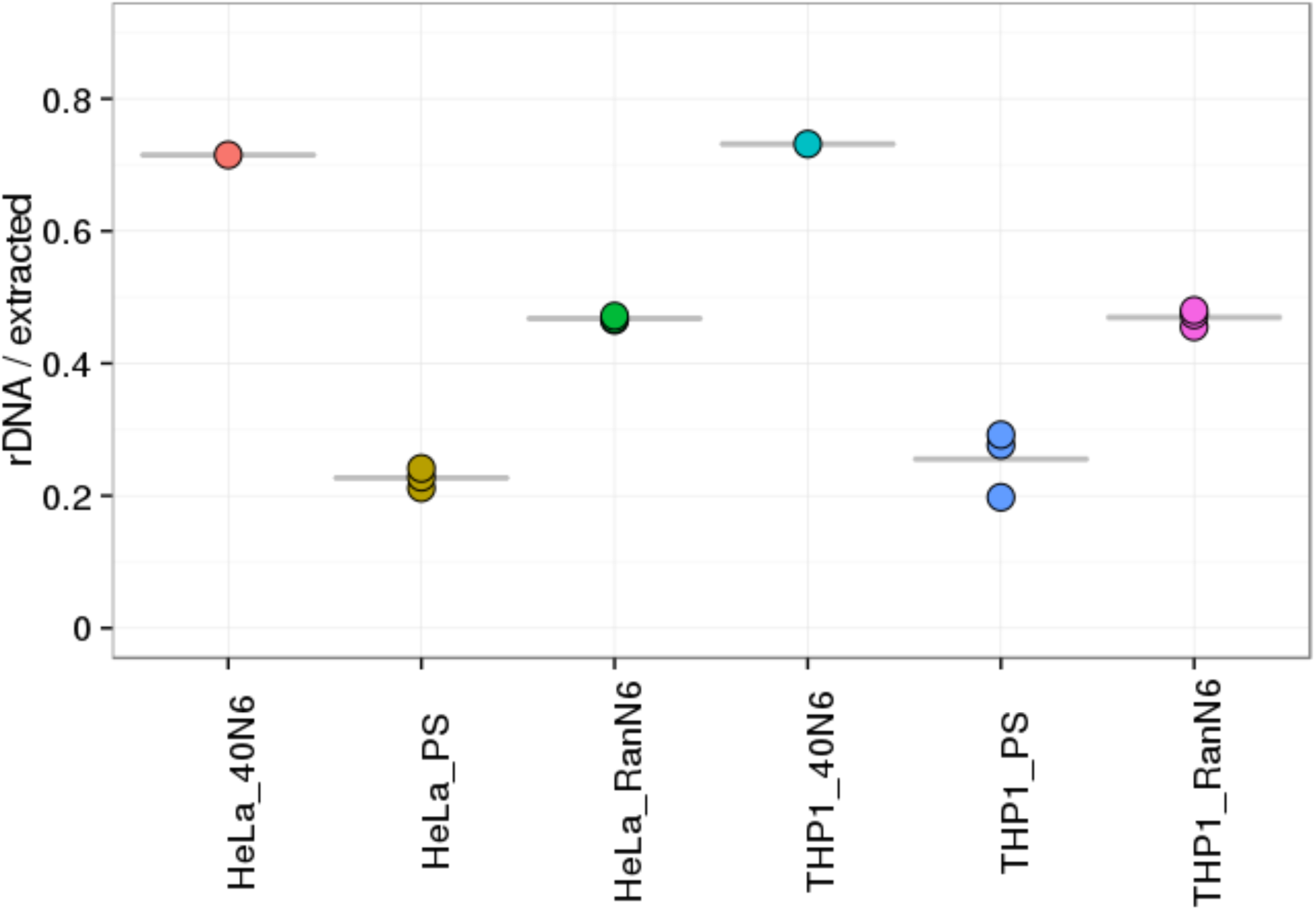

#### Numbers of genes: percentage

~~~
genes_percentage <- libs[,c(‘samplename’, ‘group’, ‘genes.sub1’)]
genes_percentage$group1 <- genes_percentage$samplename
genes_percentage$group1 <- sub(‘_.$’, ‘’, genes_percentage$group1)
genes_percentage$group1 <- factor(genes_percentage$group1)
genes_percentage <- tapply(genes_percentage$genes.sub1, genes_percentage$grou
p1, mean)
genes_percentage <- sapply(
    c(“NC12_HeLa”, “NC12_THP1”, “NC17_HeLa”, “NC17_THP1”, “NCki_HeLa”, “NCki_TH P1”),
**function**(experiment) genes_percentage[grep(experiment, names(genes_percenta ge))] / genes_percentage[paste0(experiment, “_RanN6”)] * 100
)
genes_percentage <- unlist(genes_percentage)
names(genes_percentage) <- sub(“.*\\.”, “”, names(genes_percentage))
~~~

~~~
genes_percentage <- data.frame(genes_percentage)
genes_percentage$group <- rownames(genes_percentage)
genes_percentage$group <- sub(‘NC.._’, ‘’, genes_percentage$group)
~~~

~~~
dotsize <- mean(genes_percentage$genes_percentage) / 110
p <- ggplot(genes_percentage, aes(x=group, y=genes_percentage)) +
stat_summary(fun.y=mean, fun.ymin=mean, fun.ymax=mean,
geom=“crossbar”, color=“gray”) +
              geom_dotplot(aes(fill=group), binaxis=‘y’, binwidth=1,
dotsize=dotsize, stackdir=‘center’) +
              theme_bw() +
      theme(axis.text.x = element_text(size=14, angle=90)) +
      theme(axis.text.y = element_text(size=14)) +
      theme(axis.title.y = element_blank()) +
      theme(axis.title.x = element_text(size=14))+
ylim(90,120) +
      ylab(“percentage of genes detected”)
p + guides(col = guide_legend(nrow = 8))
~~~

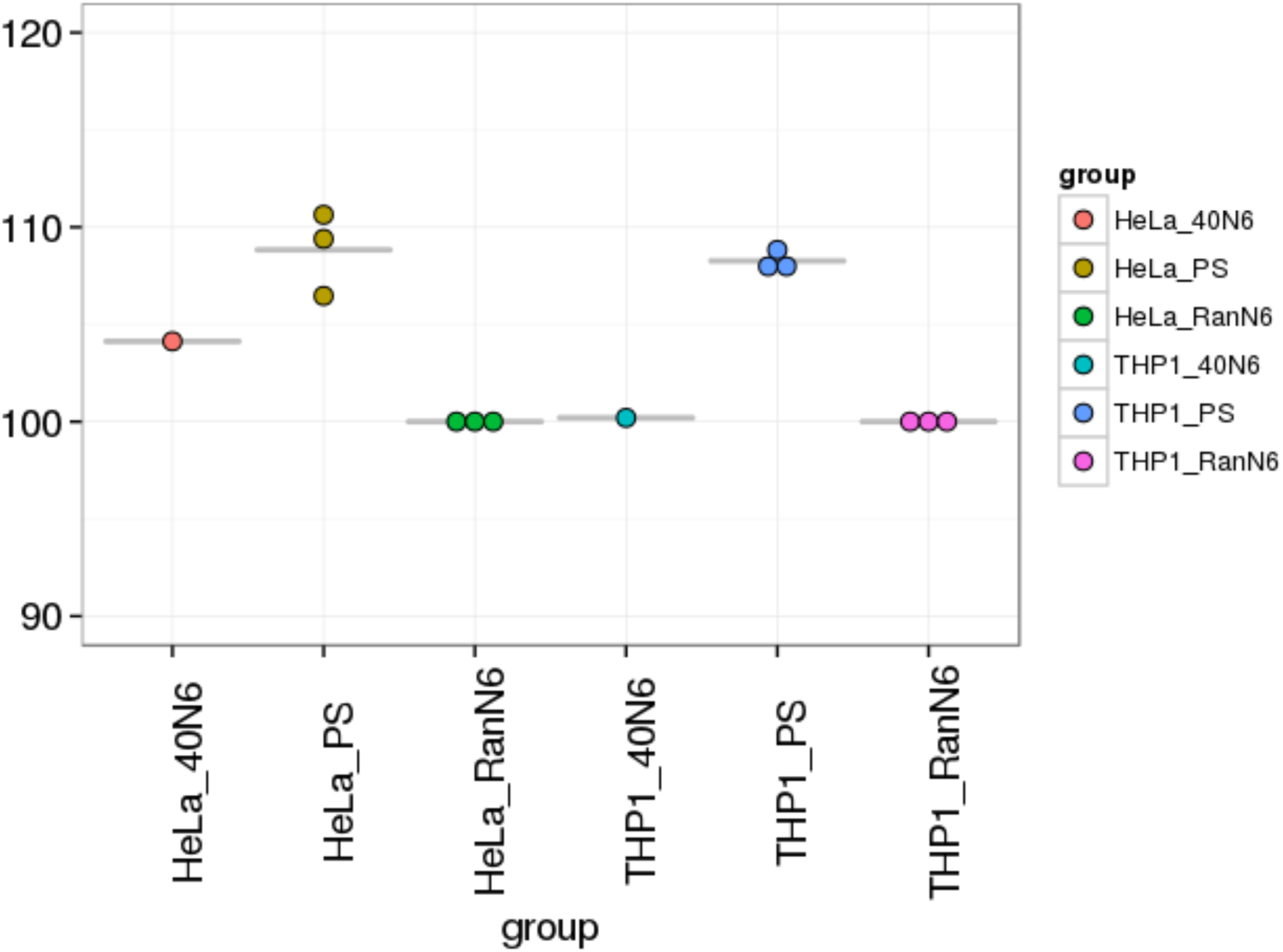

#### Transcriptome analysis

Load the data

~~~
g2_NC12 <- read.table(‘g2.sub1_NC12_1.txt’, sep=“\t”, head=T)
g2_NC17 <- read.table(‘g2.sub1_NC17_1.txt’, sep=“\t”, head=T)
g2_NCki <- read.table(‘g2.sub1_NCki_1.txt’, sep=“\t”, head=T)
~~~

Create a new table

~~~
g2 <- merge(g2_NC12, g2_NC17, by=‘row.names’, all=T)
rownames(g2) <- g2$Row.names
g2 <- g2[,-1]
g2 <- merge(g2,g2_NCki, by=‘row.names’, all=T)
rownames(g2) <- g2$Row.names
g2 <- g2[,-1]
g2[is.na(g2)] <- 0
g2b <- g2[-1,]
~~~

~~~
RanN6_HeLa = c(‘NC12.HeLa_RanN6_A’, ‘NC12.HeLa_RanN6_B’, ‘NC12.HeLa_RanN6_C’
  , ‘NC17_HeLa_RanN6_A’, ‘NC17_HeLa_RanN6_B’, ‘NC17_HeLa_RanN6_C’
    , ‘NCki_HeLa_RanN6_A’,     ‘NCki_HeLa_RanN6_B’,     ‘NCki_HeLa_RanN6_C’)
PS_HeLa = c( ‘NC12.HeLa_PS_A’,     ‘NC12.HeLa_PS_B’,     ‘NC12.HeLa_PS_C’
     , ‘NC17_HeLa_PS_A’,     ‘NC17_HeLa_PS_B’,     ‘NC17_HeLa_PS_C’
   , ‘NCki_HeLa_PS_A’,     ‘NCki_HeLa_PS_B’,     ‘NCki_HeLa_PS_C’)
RanN6_THP1 = c( ‘NC12.THP1_RanN6_A’, ‘NC12.THP1_RanN6_B’, ‘NC12.THP1_RanN6_C’
     , ‘NC17_THP1_RanN6_A’,     ‘NC17_THP1_RanN6_B’,     ‘NC17_THP1_RanN6_C’
     , ‘NCki_THP1_RanN6_A’,     ‘NCki_THP1_RanN6_B’,     ‘NCki_THP1_RanN6_C’)
PS_THP1 = c( ‘NC12.THP1_PS_A’,     ‘NC12.THP1_PS_B’,     ‘NC12.THP1_PS_C’
     , ‘NC17_THP1_PS_A’,     ‘NC17_THP1_PS_B’,     ‘NC17_THP1_PS_C’
     , ‘NCki_THP1_PS_A’,     ‘NCki_THP1_PS_B’,     ‘NCki_THP1_PS_C’)
~~~

~~~
mx <- function(DATA)
   {data.frame( HeLa_RanN6 = rowMeans(DATA[,RanN6_HeLa])
                      , HeLa_pseudoRan = rowMeans(DATA[,PS_HeLa])
                      , THP1_RanN6       = rowMeans(DATA[,RanN6_THP1])
                      , THP1_pseudoRan = rowMeans(DATA[,PS_THP1]))}
m2 <- mx(g2)
write.table(m2, “m2.txt”, sep = “\t”, quote = FALSE)
~~~

~~~
panel.cor <- function(x, y, digits=2, prefix=““, cex.cor, …)
{
usr <- par(“usr”); on.exit(par(usr))
par(usr =c(0, 1, 0, 1))
r <- abs(cor(x, y))
txt <- format(c(r, 0.123456789), digits=digits)[1]
txt <- paste(prefix, txt, sep=“”)
**if**(missing(cex.cor)) cex.cor <- 0.8/strwidth(txt)
text(0.5, 0.5, txt, cex = cex.cor * r)
}
pointsUnique <- function(x,y,…)
    points(unique(data.frame(x,y)),…)
pairPanel <- function(dataframe, title)
   pairs( dataframe
            , lower.panel=panel.cor
            , upper.panel=pointsUnique
            , main=title
            , pch=‘.’, cex=4)
~~~

~~~
pairPanel(log(m2+1), ‘pseudo-random primers’)
~~~

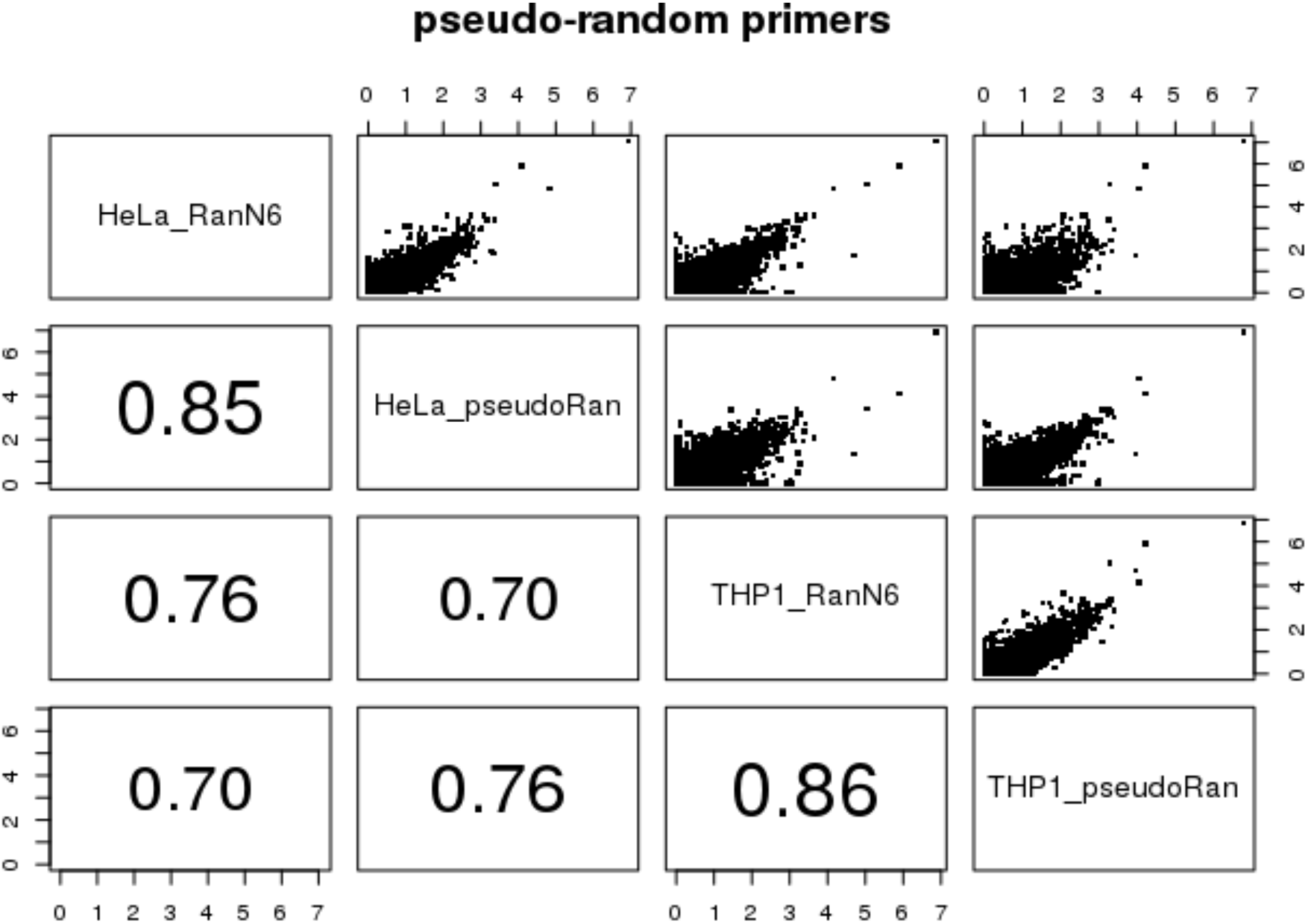

~~~
plotFoldChange <- function (DATA, COL, MAX, FUN=points, …) {
   with( DATA[rowSums(DATA) > MAX,] +1
          , FUN( HeLa_RanN6         / THP1_RanN6
                     , HeLa_pseudoRan  / THP1_pseudoRan
                     , col=COL
                     , pch=‘.’
                     , cex=5
                     , … ))
}
~~~

~~~
plotFoldChangeGrays <- function (DATA, TITLE, xlab=“Standard N6 random primer
s”, ylab=“Pseudo-random primers” ) {
   plotFoldChange( DATA,’gray90’, 0
                              , plot, log=‘xy’, main=TITLE
                              , xlab=xlab, ylab=ylab)
   plotFoldChange(DATA, ‘gray70’, 10)
   plotFoldChange(DATA, ‘gray50’, 20)
   plotFoldChange(DATA, ‘gray30’, 30)
   plotFoldChange(DATA, ‘gray10’, 40)
   legend( ‘bottomright’
              , legend=c(0, 10, 20, 30, 40)
              , col=c(‘gray90’, ‘gray70’, ‘gray50’, ‘gray30’, ‘gray10’)
              , pch=16, title=‘min expr.’)
^}^
~~~

~~~
u2 <- unique(m2)
~~~

Draw graphs

~~~
plotFoldChangeGrays(u2, “HeLa - THP-1 fold changes”)
~~~

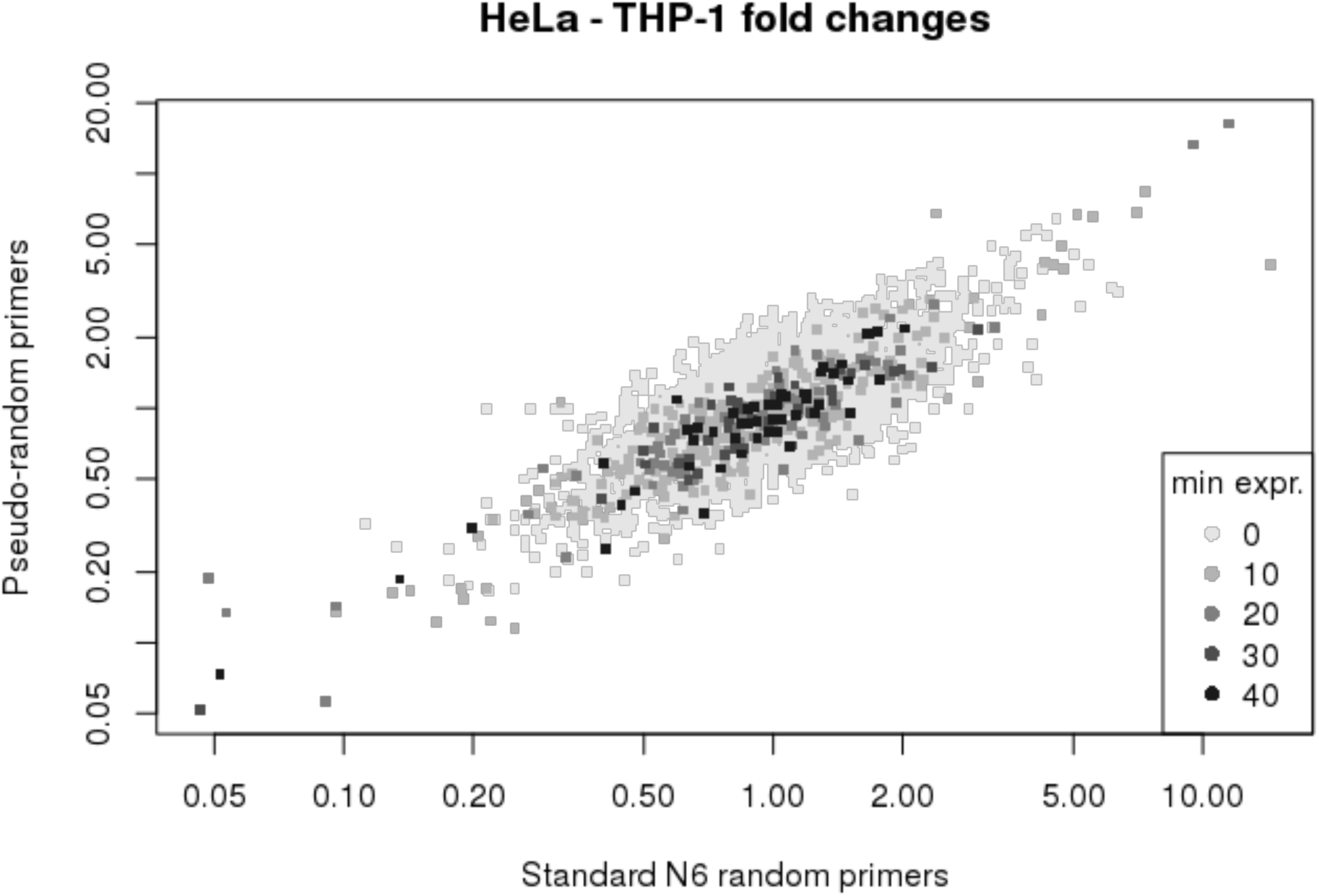

~~~
plotFoldChange( u2, ‘gray10’, 0
                          , plot, log=‘xy’, main=“HeLa - THP-1 fold changes”
                          , xlab=“Standard N6 random primers”, ylab=“Pseudo-random primer
s”)
~~~

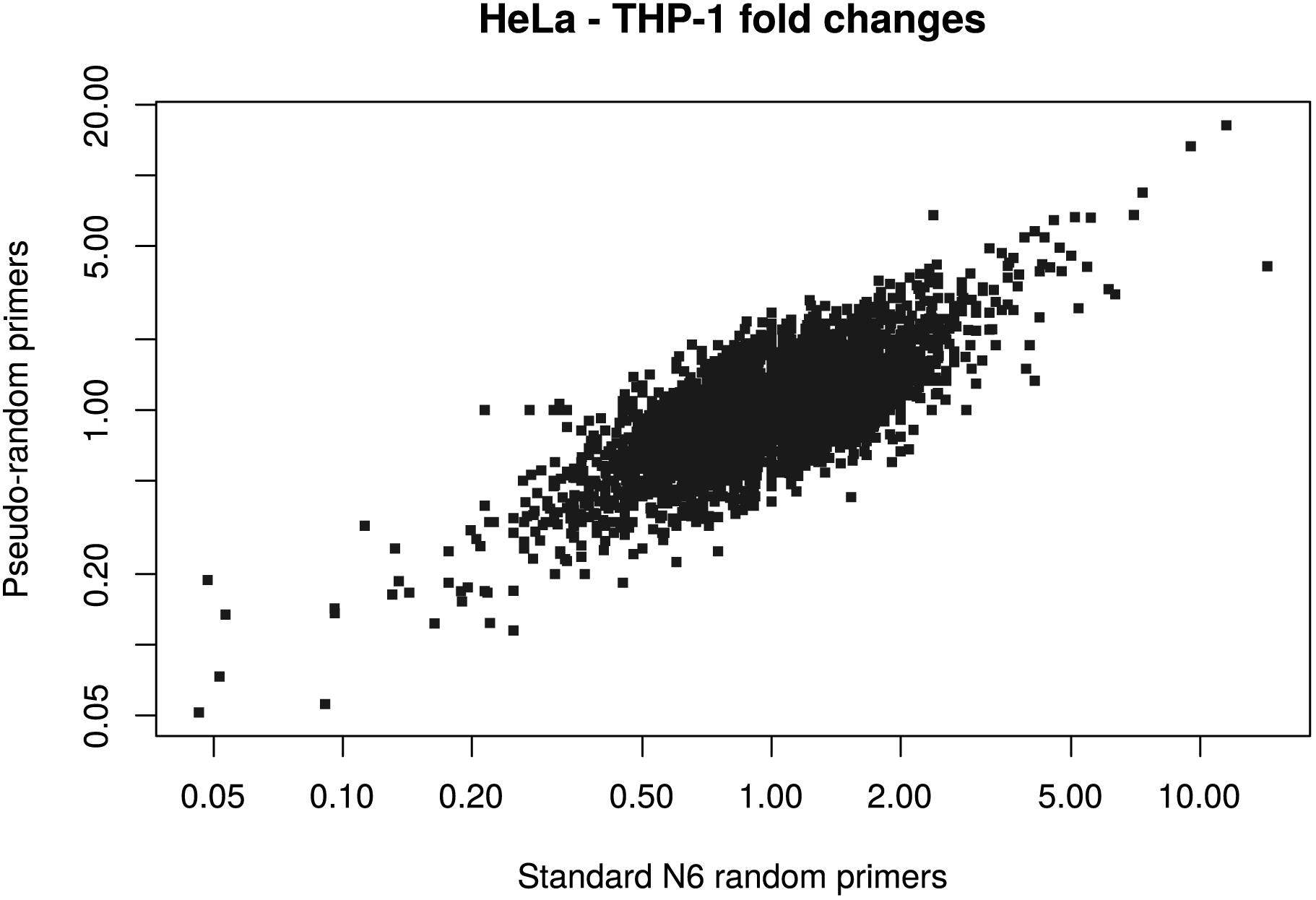

## Statistic tests

### statistic tests about sequences coming from ribosomal RNA

Regarding the PS and RanN6 set, we use a paired t.test as the the results come from 3 independents experiments.

~~~
rDNA <- read.table(‘rDNA.csv’, sep=“,”, head=T)
~~~

~~~
## Warning in read.table(“rDNA.csv”, sep = “,”, head = T): incomplete final l
ine found by readTableHeader on ‘rDNA.csv’
~~~

~~~
rDNA
~~~

~~~
##    experiments HeLa_40N6    HeLa_PS HeLa_RanN6    THP1_40N6 THP1_PS THP1_RanN 6
## 1          NC120.7149618   0.2279513   0.4715841   0.7314974   0.2765081   0.480080 5
## 2          NC17          NA 0.2413637   0.4651863            NA 0.2921605   0.473343 9
## 3          Ncki            NA 0.2116392   0.4669319            NA 0.1976666   0.455164 1
~~~

~~~
t.test(rDNA$HeLa_PS, rDNA$HeLa_RanN6, paired = T)
~~~

~~~
##
## Paired t-test
##
## data: rDNA$HeLa_PS and rDNA$HeLa_RanN6
## t = -26.2275, df = 2, p-value = 0.001451
## alternative hypothesis: true difference in means is not equal to 0
## 95 percent confidence interval:
##   -0.2804387 -0.2013934
## sample estimates:
## mean of the differences
##                       -0.2409161
~~~

~~~
t.test(rDNA$THP1_PS, rDNA$THP1_RanN6, paired = T)
~~~

~~~
##
##   Paired t-test
##
## data:   rDNA$THP1_PS and rDNA$THP1_RanN6
## t = -9.4525, df = 2, p-value = 0.01101
## alternative hypothesis: true difference in means is not equal to 0
## 95 percent confidence interval:
##   -0.3115324 -0.1166364
## sample estimates:
## mean of the differences
##                      -0.2140844
~~~

~~~
rDNA_40N6 <- read.table(‘rDNA_4 0N6.csv’, sep=“,”, head=T)
~~~

~~~
## Warning in read.table(‘‘rDNA_40N6.csv”, sep = “,”, head = T): incomplete fi
nal line found by readTableHeader on
## ‘rDNA 4 0N6.csv’
~~~

~~~
rDNA 40N6
~~~

~~~
##   experiments HeLa_40N6   HeLa_PS HeLa_RanN6 THP1_40N6   THP1_PS THP1_RanN 6
## 1      NC12_A 0.6971786 0.2241748   0.4644208 0.6973555 0.2642123 0.451782 0
## 2      NC12_B 0.72099520.2349407   0.4991978 0.7501979 0.2956299 0.518855 2
## 3      NC12_C 0.7267116 0.2247385   0.4511338 0.7469388 0.2696820 0.469604 3
~~~

Regarding the 40N6 set, we can not use the paired test as only one experiment has been performed. Thus, we use the 3 replicats of 1 experiment.

~~~
t.test(rDNA_4 0N6$HeLa_RanN6, rDNA_40N6$HeLa_4 0N6)
~~~

~~~
##
##   Welch Two Sample t-test
##
## data: rDNA_40N6$HeLa_RanN6 and rDNA_40N6$HeLa_40N6
## t = -14.363, df = 3.375, p-value = 0.0003821
## alternative hypothesis: true difference in means is not equal to 0
## 95 percent confidence interval:
##   -0.2940671 -0.1926882
## sample estimates:
## mean of x mean of y
## 0.4715841 0.7149618
~~~

~~~
t.test(rDNA_40N6$HeLa_PS, rDNA_40N6$HeLa_40N6)
~~~

~~~
##
## Welch Two Sample t-test
##
## data: rDNA_40N6$HeLa_PS and rDNA_40N6$HeLa_40N6
## t = -50.2252, df = 2.586, p-value = 6.162e-05
## alternative hypothesis: true difference in means is not equal to 0
## 95 percent confidence interval:
##    -0.5208657 -0.4531552
## sample estimates:
## mean of x mean of y
## 0.2279513 0.7149618
~~~

~~~
t.test(rDNA_4 0N6$THP1_RanN6, rDNA_40N6$THP1_40N6)
~~~

~~~
##
## Welch Two Sample t-test
##
## data: rDNA_40N6$THP1_RanN6 and rDNA_40N6$THP1_40N6
## t = -9.5392, df = 3.902, p-value = 0.0007598
## alternative hypothesis: true difference in means is not equal to 0
## 95 percent confidence interval:
##   -0.3253230 -0.1775109
## sample estimates:
## mean of x mean of y
## 0.4800805 0.7314974
~~~

~~~
t.test(rDNA_4 0N6$THP1_PS, rDNA_40N6$THP1_40N6)
~~~

~~~
##
## Welch Two Sample t-test
##
## data: rDNA_40N6$THP1_PS and rDNA_40N6$THP1_40N6
## t = -23.1521, df = 3.165, p-value = 0.0001224
## alternative hypothesis: true difference in means is not equal to 0
## 95 percent confidence interval:
##   -0.5157285 -0.3942503
## sample estimates:
## mean of x mean of y
## 0.2765081 0.7314974
~~~

### statistic tests about sequences coming from artefacts

Regarding the PS and RanN6 set, we use a paired t.test as the the results come from 3 independents experiments.

~~~
artefact <- read.table(‘artefacts.csv’, sep=“,”, head=T)
~~~

~~~
## Warning in read.table(“artefacts.csv”, sep = “,”, head = T): incomplete fi
nal line found by readTableHeader on
## ‘artefacts.csv’
~~~

~~~
artefact
~~~

~~~
##    experiments    HeLa_40N6      HeLa_PS HeLa_RanN6    THP1_40N6      THP1_PS THP1_
RanN6
## 1            NC12 0.01044525 0.01139030 0.08445126 0.01031001 0.01440960 0.088
87914
## 2            NC17                NA 0.02860542 0.02765798             NA 0.02274970 0.075
05295
## 3            Ncki                NA 0.02998795 0.07369473               NA 0.02369741 0.061
45156
~~~

~~~
t.test(artefact$HeLa_PS, artefact$HeLa_RanN6, paired = T)
~~~

~~~
##
## Paired t-test
##
## data: artefact$HeLa_PS and artefact$HeLa_RanN6
## t = -1.7943, df = 2, p-value = 0.2146
## alternative hypothesis: true difference in means is not equal to 0
## 95 percent confidence interval:
##   -0.13118278 0.05396924
## sample estimates:
## mean of the differences
##                      -0.03860677
~~~

~~~
t.test(artefact$THP1_PS, artefact$THP1_RanN6, paired = T)
~~~

~~~
##
## Paired t-test
##
## data: artefact$THP1_PS and artefact$THP1_RanN6
## t = -5.1377, df = 2, p-value = 0.03586
## alternative hypothesis: true difference in means is not equal to 0
## 95 percent confidence interval:
##   -0.100771328 -0.008913305
## sample estimates:
## mean of the differences
##                      -0.05484232
~~~

Regarding the 40N6 set, we can not use the paired test as only one experiment has been performed. Thus, we use the 3 replicats of 1 experiment.

~~~
artefact_40N6 <- read.table(‘artefact_40N6.csv’, sep=“,”, head=T)
~~~

~~~
## Warning in read.table(“artefact_40N6.csv”, sep = head = T): incomplet
e final line found by readTableHeader on
## ‘artefact 40N6.csv’
~~~

~~~
artefact_40N6
~~~

~~~
##    experiments    HeLa_40N6    HeLa_PS HeLa_RanN6    THP1_40N6      THP1_PS TH
P1_RanN6
## 1      NC12_A 0.014282432 0.014614436 0.08701817 0.01415881 0.019349645 0.
13564309
## 2      NC12_B 0.005714088 0.007657406 0.05647798 0.00548908 0.008236931 0.
03813052
## 3      NC12_C 0.011339241 0.011899050 0.10985765 0.01128215 0.015642227 0.
09286382
~~~

~~~
t.test(artefact_40N6$HeLa_RanN6, artefact_40N6$HeLa_40N6)
~~~

~~~
##
## Welch Two Sample t-test
##
## data: artefact_40N6$HeLa_RanN6 and artefact_40N6$HeLa_40N6
## t = 4.7241, df = 2.106, p-value = 0.03798
## alternative hypothesis: true difference in means is not equal to 0 ## 95 percent confidence interval:
##   0.009740573 0.138271448
## sample estimates:
##   mean of x mean of y
## 0.08445126 0.01044525
~~~

~~~
t.test(artefact_40N6$HeLa_PS, artefact_40N6$HeLa_40N6)
~~~

~~~
##
## Welch Two Sample t-test
##
## data: artefact_40N6$THP1_RanN6 and artefact_40N6$THP1_40N6
## t = 2.7729, df = 2.033, p-value = 0.1073
## alternative hypothesis: true difference in means is not equal to 0
## 95 percent confidence interval:
## -0.04148839 0.19862666
## sample estimates:
## mean of x mean of y
## 0.08887914 0.01031001
~~~

~~~
t.test(artefact_40N6$THP1_PS, artefact_40N6$THP1_40N6)
~~~

~~~
##
##   Welch Two Sample t-test
##
## data: artefact_40N6$THP1_RanN6 and artefact_40N6$THP1_40N6
## t = 2.7729, df = 2.033, p-value = 0.1073
## alternative hypothesis: true difference in means is not equal to 0
## 95 percent confidence interval:
##   -0.04148839 0.19862666
## sample estimates:
##   mean of x mean of y
## 0.08887914 0.01031001
~~~

~~~
t.test(artefact_40N6$THP1_PS, artefact_40N6$THP1_40N6)
~~~

~~~
##
## Welch Two Sample t-test
##
## data: artefact_40N6$THP1_PS and artefact_40N6$THP1_40N6
## t = 0.9893, df = 3.777, p-value = 0.3816
## alternative hypothesis: true difference in means is not equal to 0
## 95 percent confidence interval:
## -0.007678089 0.015877266
## sample estimates:
## mean of x mean of y
## 0.01440960 0.01031001
~~~

### statistic tests about the numbers of genes detected

Regarding the PS and RanN6 set, we use a paired t.test as the the results come from 3 independents experiments.

~~~
genes <- read.table(‘genes.csv’, sep=“,”, head=T)
~~~

~~~
## Warning in read.table(“genes.csv”, sep = “,”, head = T): incomplete final line found by readTableHeader on ‘genes.csv’
~~~

~~~
genes
~~~

~~~
##    experiments HeLa_40N6   HeLa_PS HeLa_RanN6   THP1_40N6 THP1_PS THP1_RanN6
## 1           NC12 104.1283 110.6335             100 100.1942 108.2811           100
## 2           NC17            NA 106.4641           100           NA 107.6821           f100
## 3           Ncki           NA 109.3965           100           NA 108.8293           100
~~~

~~~
t.test(genes$HeLa_PS, genes$HeLa_RanN6, paired = T)
~~~

~~~
##
## Paired t-test
##
## data: genes$HeLa_PS and genes$HeLa_RanN6
## t = 7.1433, df = 2, p-value = 0.01904
## alternative hypothesis: true difference in means is not equal to 0
## 95 percent confidence interval:
##     3.511913 14.150874
## sample estimates:
## mean of the differences
##                         8.831394
~~~

~~~
t.test(genes$THP1_PS, genes$THP1_RanN6, paired = T)
~~~

~~~
##
## Paired t-test
##
## data: genes$THP1_PS and genes$THP1_RanN6
## t = 24.9454, df = 2, p-value = 0.001603
## alternative hypothesis: true difference in means is not equal to 0
## 95 percent confidence interval:
## 6.83873 9.68958
## sample estimates:
## mean of the differences
##                         8.264155
~~~

Regarding the 40N6 set, we can not use the paired test as only one experiment has been performed. Thus, we use the 3 replicats of 1 experiment.

~~~
genes_40N6 <- read.table(‘genes_40N6.csv’, sep=“,”, head=T)
~~~

~~~
## Warning in read.table(“genes_40N6.csv”, sep = “,”, head = T): incomplete f
inal line found by readTableHeader on
## ‘genes_40N6.csv’
~~~

~~~
genes_4 0N6
~~~

\## experiments HeLa_40N6 HeLa_PS HeLa_RanN6 THP1_40N6 THP1_PS THP1_RanN6

~~~
## 1           NC12_A   102.3233 108.8375      96.80100   94.68527 106.6037   96.35384
## 2           NC12_B   103.3688 110.0974   103.87812 101.91578 106.1005   102.55143
## 3           NC12_C   106.6929 112.9658   99.32088 103.98164 112.1391   101.09473
~~~

~~~
t.test(genes_40N6$HeLa_RanN6, genes_40N6$HeLa_40N6)
~~~

~~~
##
## Welch Two Sample t-test
##
## data: genes_40N6$HeLa_RanN6 and genes_40N6$HeLa_40N6
## t = -1.682, df = 3.391, p-value = 0.1806
## alternative hypothesis: true difference in means is not equal to 0
## 95 percent confidence interval:
##   -11.454223     3.197589
## sample estimates:
## mean of x mean of y
##   100.0000   104.1283
~~~

~~~
t.test(genes_40N6$HeLa_PS, genes_40N6$HeLa_40N6)
~~~

~~~
##
## Welch Two Sample t-test
##
## data: genes_40N6$HeLa_PS and genes_40N6$HeLa_40N6
## t = 3.621, df = 3.977, p-value = 0.02255
## alternative hypothesis: true difference in means is not equal to 0
## 95 percent confidence interval:
##   1.506129 11.504326
## sample estimates:
## mean of x mean of y
##   110.6335 104.1283
~~~

~~~
t.test(genes_40N6$THP1_RanN6, genes_40N6$THP1_40N6)
~~~

~~~
##
## Welch Two Sample t-test
##
## data: genes_40N6$THP1_RanN6 and genes_40N6$THP1_40N6
## t = -0.0574, df = 3.476, p-value = 0.9574
## alternative hypothesis: true difference in means is not equal to 0
## 95 percent confidence interval:
##   -10.171798 9.783346
## sample estimates:
## mean of x mean of y
##   100.0000   100.1942
~~~

~~~
t.test(genes_40N6$THP1_PS, genes_40N6$THP1_40N6)
~~~

~~~
##
## Welch Two Sample t-test
##
## data: genes_40N6$THP1_PS and genes_40N6$THP1_40N6
## t = 2.3657, df = 3.542, p-value = 0.08558
## alternative hypothesis: true difference in means is not equal to 0
## 95 percent confidence interval:
## -1.90779 18.08153
## sample estimates:
## mean of x mean of y
## 108.2811 100.1942
~~~

## Targeted reduction of Hemoglobin cDNAs

### Configuration

~~~
**library**(plyr)
exportInEnv <- **function**(X) {
    Name    <- X
    Value    <- get(X)
    .Internal(Sys.setenv(Name, Value))
    cat( paste0(“export “, paste(Name, Value, sep=‘=‘), “\n”))
^}^
LIBRARY                  <- ‘NC22b’
MOIRAI_USER         <- ‘nanoCAGE2’
MOIRAI_PROJECT  <- ‘Arnaud’
GROUP_SHARED    <- ‘/osc-fs_home/scratch/gmtu’
WORKDIR                <- ‘.’
GENE_SYMBOLS <- paste(GROUP_SHARED, ‘annotation/homo_sapiens/gencode-14/gen code.v14.annotation.genes.bed’, sep=‘/’)
ANNOTATION <- paste(GROUP_SHARED, ‘annotation/homo_sapiens/100 712hg19/100
712hg19’, sep=‘/’)
PROCESSED_DATA <- dirname( system( paste( ‘ls -d /osc-fs_home/scratch/moirai/
                                                                     , MOIRAI_USER
                                                                     , ‘/project/’
                                                                     , MOIRAI_PROJECT
                                                                     , ‘/’
                                                                     , LIBRARY
                                                                     , ‘*/Moirai.config’
                                                                     , sep=‘’)
                                                           , intern=TRUE)[1])
l_ply( c(“LIBRARY”, “MOIRAI_USER”, “MOIRAI_PROJECT”, “GROUP_SHARED”
              , “WORKDIR”, “GENE_SYMBOLS”, “ANNOTATION”, “PROCESSED_DATA”)
          , exportInEnv)
~~~

~~~
export LIBRARY=NC22b
export MOIRAI_USER=nanoCAGE2
export MOIRAI_PROJECT=Arnaud
export GROUP_SHARED=/osc-fs_home/scratch/gmtu
export WORKDIR=.
export GENE_SYMBOLS=/osc-fs_home/scratch/gmtu/annotation/homo_sapiens/gencode-14/gencode.v14.annotation.genes.bed
export ANNOTATION=/osc-fs_home/scratch/gmtu/annotation/homo_sapiens/100 712hg1 9/100712hg19
export PROCESSED_DATA=/osc-fs_home/scratch/moirai/nanoCAGE2/project/Arnaud/NC 22b.CAGEscan short-reads.20150625152335
~~~

Moirai URL: http://moirai.gsc.riken.jp/osc-fs_home/scratch/moirai/nanoCAGE2/project/Arnaud/NC22b.CAGEscan_short-reads.20150625152335/NC22b.CAGEscan_short-reads.20150625152335.html (http://moirai.gsc.riken.jp/osc-fs_home/scratch/moirai/nanoCAGE2/project/Arnaud/NC22b.CAGEscan_short-reads.20150625152335/NC22b.CAGEscan_short-reads.20150625152335.html)

## Count the reads

~~~
awk ‘/raw/ {print $3}’ $PROCESSED_DATA/text/summary.txt |
/usr/lib/filo/stats |
grep ‘Sum’ |
cut -f2 -d’:’ |
tr -d ‘[:space:]’ |
xargs -0 /usr/bin/printf “ # %’d\n”
grep raw $PROCESSED_DATA/text/summary.txt
~~~

~~~
## # 2999748
## NC22b.ACAGTG.R1 raw 181519
## NC22b.ATCACG.R1 raw 211629
## NC22b.CGATGT.R1 raw 82773
## NC22b.GCCAAT.R1 raw 170418
## NC22b.TGACCA.R1 raw 58532
## NC22b.TTAGGC.R1 raw 188190
## NC22b.Undetermined.R1      raw 2106687
~~~

## Analysis with R

### Configuration

~~~
**library**(oscR) # *See https://github.com/charles-plessy/oscR for oscR.*
**if** (compareVersion(sessionInfo()$otherPkgs$oscR$Version,’0.1.1’) < 0) stop(‘O
utdated version of oscR.’)
**library**(smallCAGEqc) *# See https://github.com/charles-plessy/smallCAGEqc for smallCAGEqc.*
**if** (compareVersion(sessionInfo()$otherPkgs$smallCAGEqc$Version,’0.6.0’) < 0)
**stop**(‘Outdated version of smallCAGEqc’)
**library**(vegan)
~~~

~~~
library(ggplot2)
~~~

### Load data

~~~
l1 <- read.osc(paste(LIBRARY, ‘l1’, ‘gz’, sep=‘.’), drop.coord=T, drop.norm=T)
l2 <- read.osc(paste(LIBRARY, ‘l2’, ‘gz’, sep=‘.’), drop.coord=T, drop.norm=T)
colnames(l1) <- sub(‘raw.NC22b.’,’’,colnames(l1))
colnames(l2) <- sub(‘raw.NC22b.’,’’,colnames(l2))
colSums(l2)
~~~

~~~
##   22_PSHb_A  22_PSHb_B  22_PSHb_C  22_RanN6_A  22_RanN6_B  22_RanN6_C
##              3786              3196              6805              17433              18864              17218
~~~

~~~
PSHb <- c(‘22_PSHb_A’, ‘22_PSHb_B’, ‘22_PSHb_C’)
RanN6     <- c(‘22_RanN6_A’, ‘22_RanN6_B’, ‘22_RanN6_C’)
~~~

### Normalization number of read per sample: libs2.sub

Libraries contain only very few reads tags. The smallest one has 3,191 counts. In order to make meaningful comparisons, all of them are subsapled to 3190 counts.

~~~
l2.sub <- t(rrarefy(t(l2),3190))
colSums(l2.sub)
~~~

~~~
##  22_PSHb_A  22_PSHb_B  22_PSHb_C  22_RanN6_A  22_RanN6_B  22_RanN6_C
##              3190              3190              3190                3190                3190                3190
~~~

### Moirai statistics

~~~
libs <- loadLogs(‘moirai’)
libs <- libs[colnames(l1),]
~~~

### Number of clusters

~~~
libs[“l2.sub”]        <- colSums(l2.sub > 0)
libs[“l2.sub.exp”]  <- rarefy(t(l2), min(colSums(l2)))
~~~

### Richness

Richness should also be calculated on the whole data.

~~~
libs[“r100.l2”] <- rarefy(t(l2),100)
t.test(data=libs, r100.l2 ~ group)
~~~

~~~
##
## Welch Two Sample t-test
##
## data: r10 0.l2 by group
## t = 13.0614, df = 3.836, p-value = 0.0002544
## alternative hypothesis: true difference in means is not equal to 0
## 95 percent confidence interval:
##     7.645323 11.863046
## sample estimates:
## mean in group PS_Hb mean in group RanN6
##                    93.44089                    83.68671
~~~

~~~
boxplot(data=libs, r100.l2 ~ group, ylim=c(80,100), las=1)
~~~

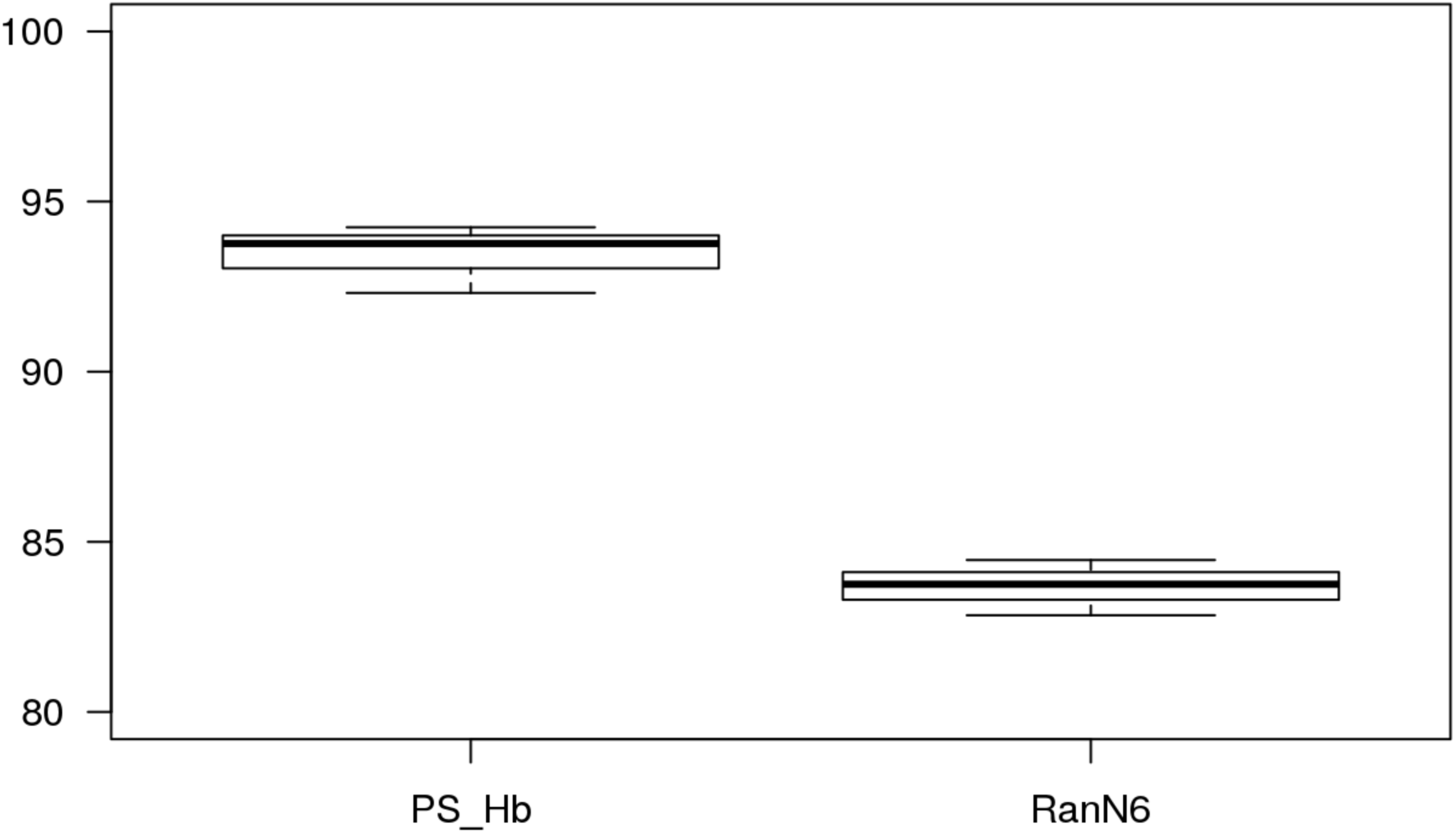

### Hierarchical annotation

~~~
annot.l2 <- read.table(paste(LIBRARY,’l2’,’annot’,sep=‘.’), head=F, col.names
=c(‘id’, ‘feature’), row.names=1)
annot.l2 <- hierarchAnnot(annot.l2)
libs <- cbind(libs, t(rowsum(l2,   annot.l2[,’class’])))
~~~

### Gene symbols used normalisation data

~~~
genesymbols <- read.table(paste(LIBRARY,’l2’,’genes’,sep=‘.’), col.names=c(“c
luster”,”symbol”), stringsAsFactors=FALSE)
rownames(genesymbols) <- genesymbols$cluster
countSymbols <- **function**(X) length(unique(genesymbols[X > 0,’symbol’]))
libs[colnames(l2.sub), “genes.sub”] <- apply(l2.sub, 2, countSymbols)
libs[colnames(l2),             “genes”] <- apply(l2,         2, countSymbols)
~~~

~~~
dotsize <- mean(libs$genes.sub) /150
par(mar=c(7,10,2,30))
p <- ggplot(libs, aes(x=group, y=genes.sub)) +
stat_summary(fun.y=mean, fun.ymin=mean, fun.ymax=mean,
geom=“crossbar”, color=“gray”) +
              geom_dotplot(aes(fill=group), binaxis=‘y’, binwidth=1,
dotsize=dotsize, stackdir=‘center’) +
              theme_bw() +
     theme(axis.text.x = element_text(size=14)) +
     theme(axis.text.y = element_text(size=14)) +
     theme(axis.title.x = element_blank())+
     theme(axis.title.y = element_text(size=14))+
ylim(1300,1600) +
   ylab(“Number of genes detected”)
p + theme(legend.position=“none”)
~~~

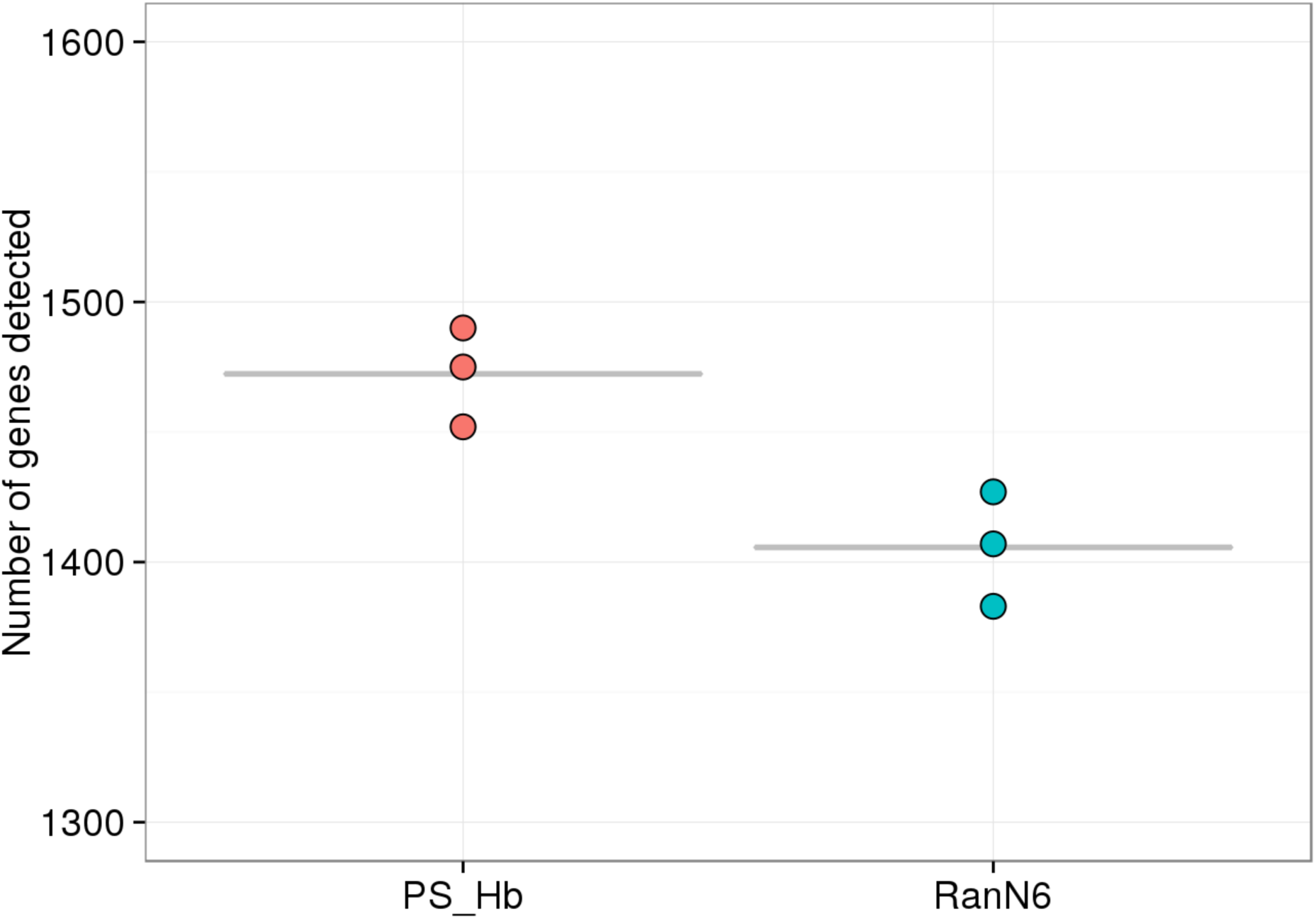

statistical analysis of gene count (with normalized data)

~~~
t.test(data=libs, genes.sub ~ group)
~~~

~~~
##
## Welch Two Sample t-test
##
## data: genes.sub by group
## t = 3.9567, df = 3.923, p-value = 0.01736
## alternative hypothesis: true difference in means is not equal to 0
## 95 percent confidence interval:
##    19.52393 113.80940
## sample estimates:
## mean in group PS_Hb mean in group RanN6
##                    1472.333                    1405.667
~~~

### Analysis of the gene expressed in different sample with different primers - normalized data (l2.sub)

~~~
l2_to_g2 <- function(l2) {
   g2 <- rowsum(l2, genesymbols$symbol)
   subset(g2, rowSums(g2) > 0)
^}^
g2.sub <- l2_to_g2(l2.sub)
g2       <- l2_to_g2(l2)
G2  <- TPM(g2)
libs$genes.r <- rarefy(t(g2), 3190)[rownames(libs)]
t.test(data=libs, genes.r ~ group)
~~~

~~~
##
## Welch Two Sample t-test
##
## data: genes.r by group
## t = 2.8877, df = 3.518, p-value = 0.05212
## alternative hypothesis: true difference in means is not equal to 0
## 95 percent confidence interval:
##    -1.227913 157.191500
## sample estimates:
## mean in group PS_Hb mean in group RanN6
##                     1491.744                     1413.763
~~~

~~~
G2mean <- function(TABLE)
     data.frame( RanN6 = TPM(rowSums(TABLE[,RanN6]))
                      , PS_Hb = TPM(rowSums(TABLE[,PSHb])))
G2.sub.mean <- G2mean(g2.sub)
G2.mean       <- G2mean(g2)
~~~

~~~
head(G2.sub.mean[order(G2.sub.mean$RanN6, decreasing=TRUE),], 30)
~~~

~~~
head(G2.sub.mean[order(G2.sub.mean$RanN6, decreasing=TRUE),], 30)
~~~

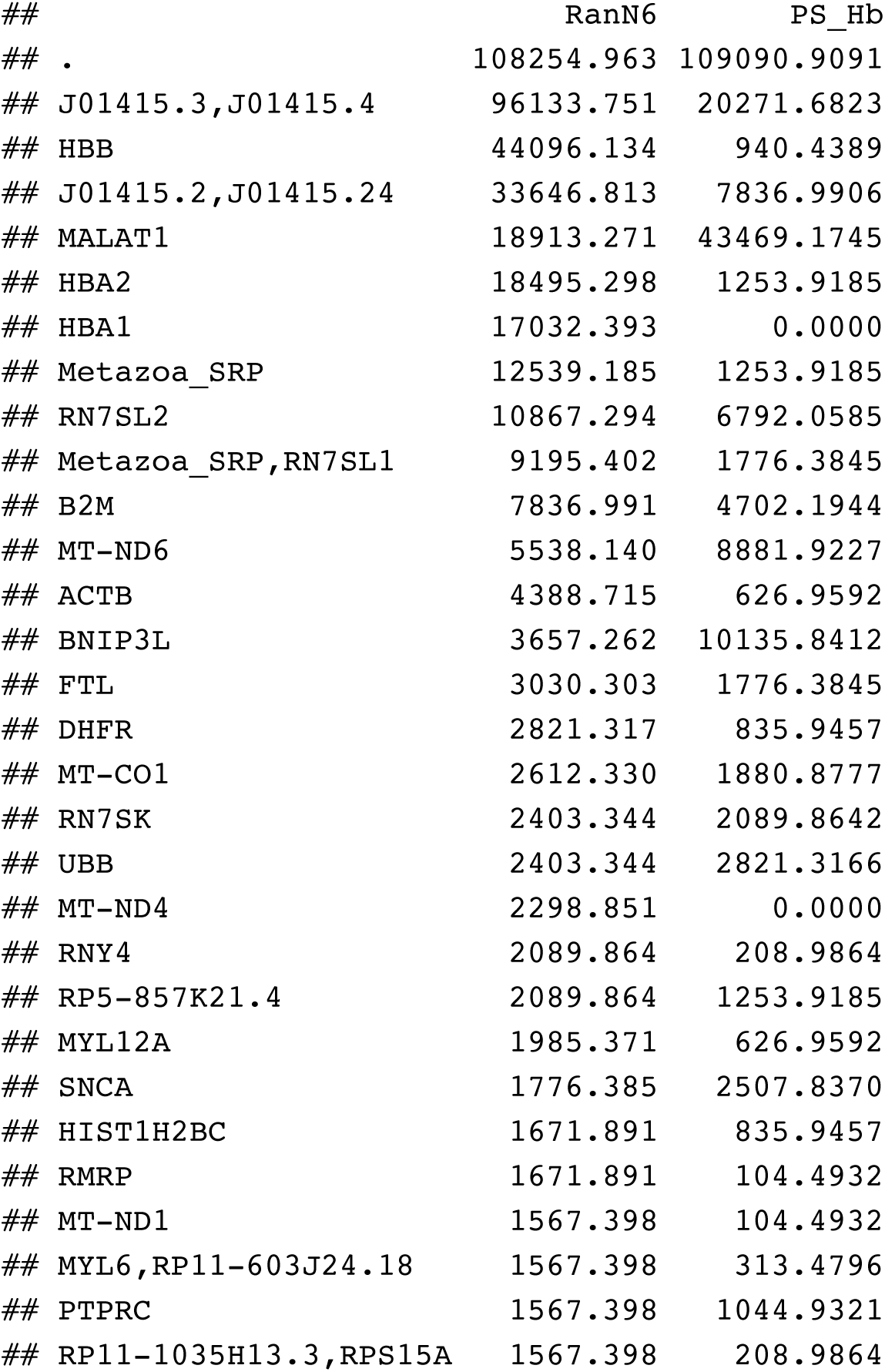

| ## | RanN6 | PS_Hb |
| --- | --- | --- |
| ## . | 108254.963 | 109090.9091 |
| ## J01415.3,J01415.4 | 96133.751 | 20271.6823 |
| ## HBB | 44096.134 | 940.4389 |
| ## J01415.2,J01415.24 | 33646.813 | 7836.9906 |
| ## MALAT1 | 18913.271 | 43469.1745 |
| ## HBA2 | 18495.298 | 1253.9185 |
| ## HBA1 | 17032.393 | 0.0000 |
| ## Metazoa_SRP | 12539.185 | 1253.9185 |
| ## RN7SL2 | 10867.294 | 6792.0585 |
| ## Metazoa_SRP,RN7SL1 | 9195.402 | 1776.3845 |
| ## B2M | 7836.991 | 4702.1944 |
| ## MT-ND6 | 5538.140 | 8881.9227 |
| ## ACTB | 4388.715 | 626.9592 |
| ## BNIP3L | 3657.262 | 10135.8412 |
| ## FTL | 3030.303 | 1776.3845 |
| ## DHFR | 2821.317 | 835.9457 |
| ## MT-CO1 | 2612.330 | 1880.8777 |
| ## RN7SK | 2403.344 | 2089.8642 |
| ## UBB | 2403.344 | 2821.3166 |
| ## MT-ND4 | 2298.851 | 0.0000 |
| ## RNY4 | 2089.864 | 208.9864 |
| ## RP5-857K21.4 | 2089.864 | 1253.9185 |
| ## MYL12A | 1985.371 | 626.9592 |
| ## SNCA | 1776.385 | 2507.8370 |
| ## HIST1H2BC | 1671.891 | 835.9457 |
| ## RMRP | 1671.891 | 104.4932 |
| ## MT-ND1 | 1567.398 | 104.4932 |
| ## MYL6,RP11-603J24.18 | 1567.398 | 313.4796 |
| ## PTPRC | 1567.398 | 1044.9321 |
| ## RP11-1035H13.3,RPS15A | 1567.398 | 208.9864 |

~~~
head(G2.sub.mean[order(G2.sub.mean$PS_Hb, decreasing=TRUE),], 30)
~~~

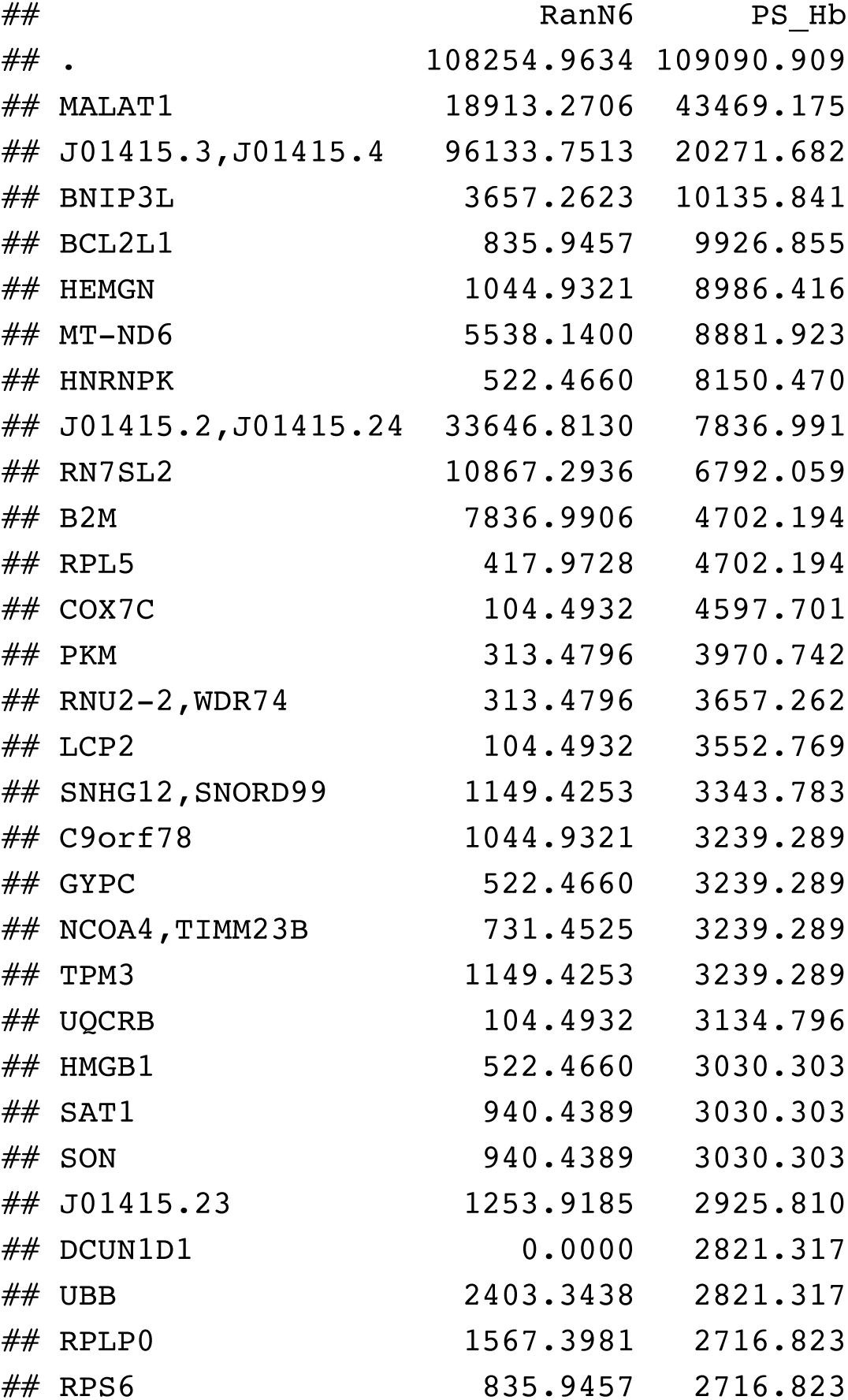

### Gene list on normalized data (table l2.sub)

~~~
RanN6_genelist.sub <- listSymbols(rownames(subset(G2.sub.mean, RanN6>0)))
PSHb_genelist.sub <- listSymbols(rownames(subset(G2.sub.mean, PS_Hb>0)))
~~~

~~~
genelist <- listSymbols(rownames(g2))
~~~

~~~
write.table(genelist, ‘NC2 2.genelist.txt’, sep = “\t”, quote = FALSE, row.names = FALSE, col.names = FALSE)
~~~

### Haemoglobin barplot

~~~
par(mar=c(2,2,2,2))
barplot(t(G2[grep(‘^HB[AB]’, rownames(g2), value=T),]), beside=T, ylab=‘Norma
lised expression value (cpm).’, col=c(“gray50”,”gray50”, “gray50”, “gray90”,
“gray90”, “gray90”))
legend(“topleft”, legend=c(“RanN6”, “PS_Hb”), fill=c(“gray90”, “gray50”))
~~~

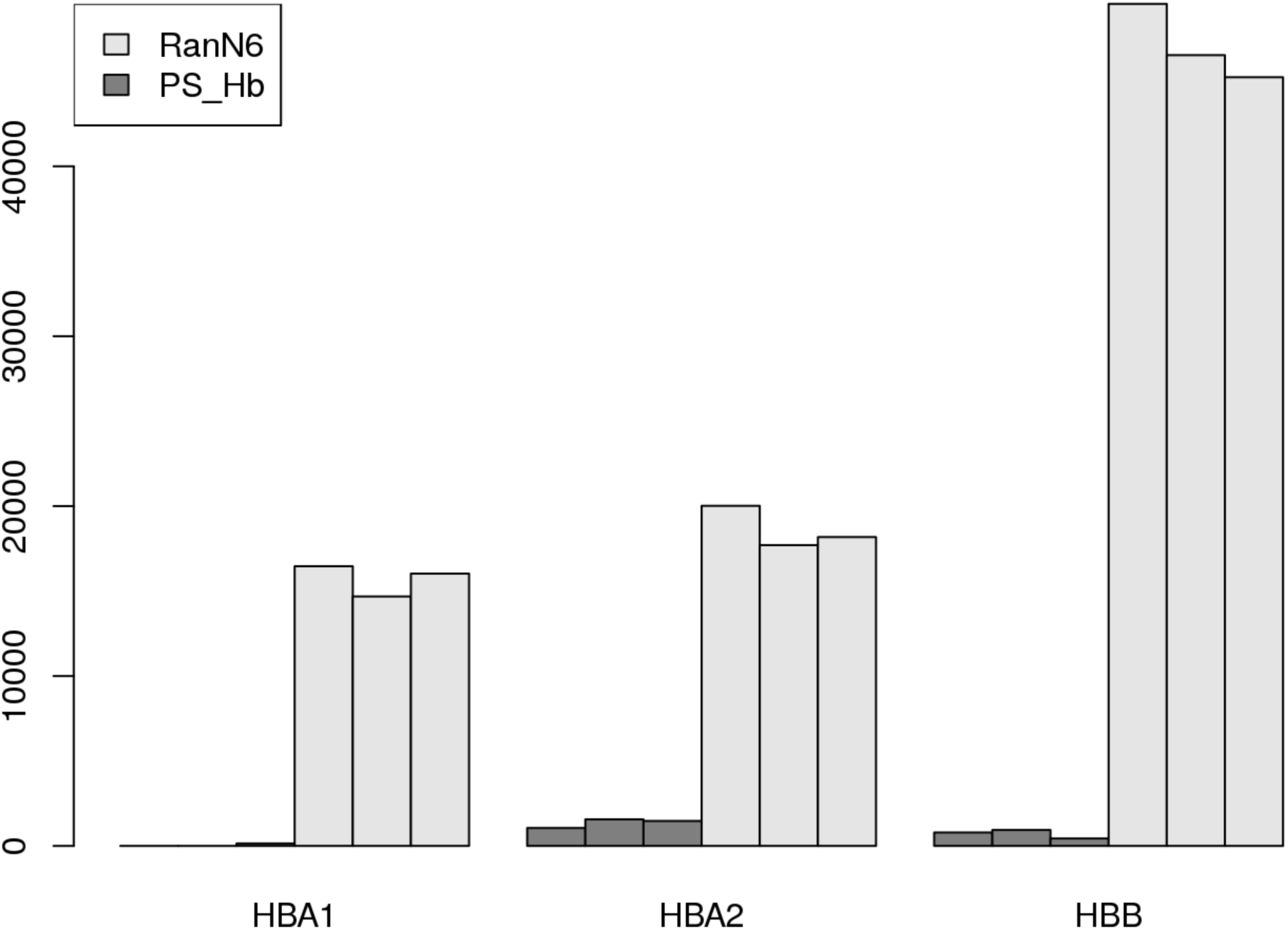

**Figure S1.**
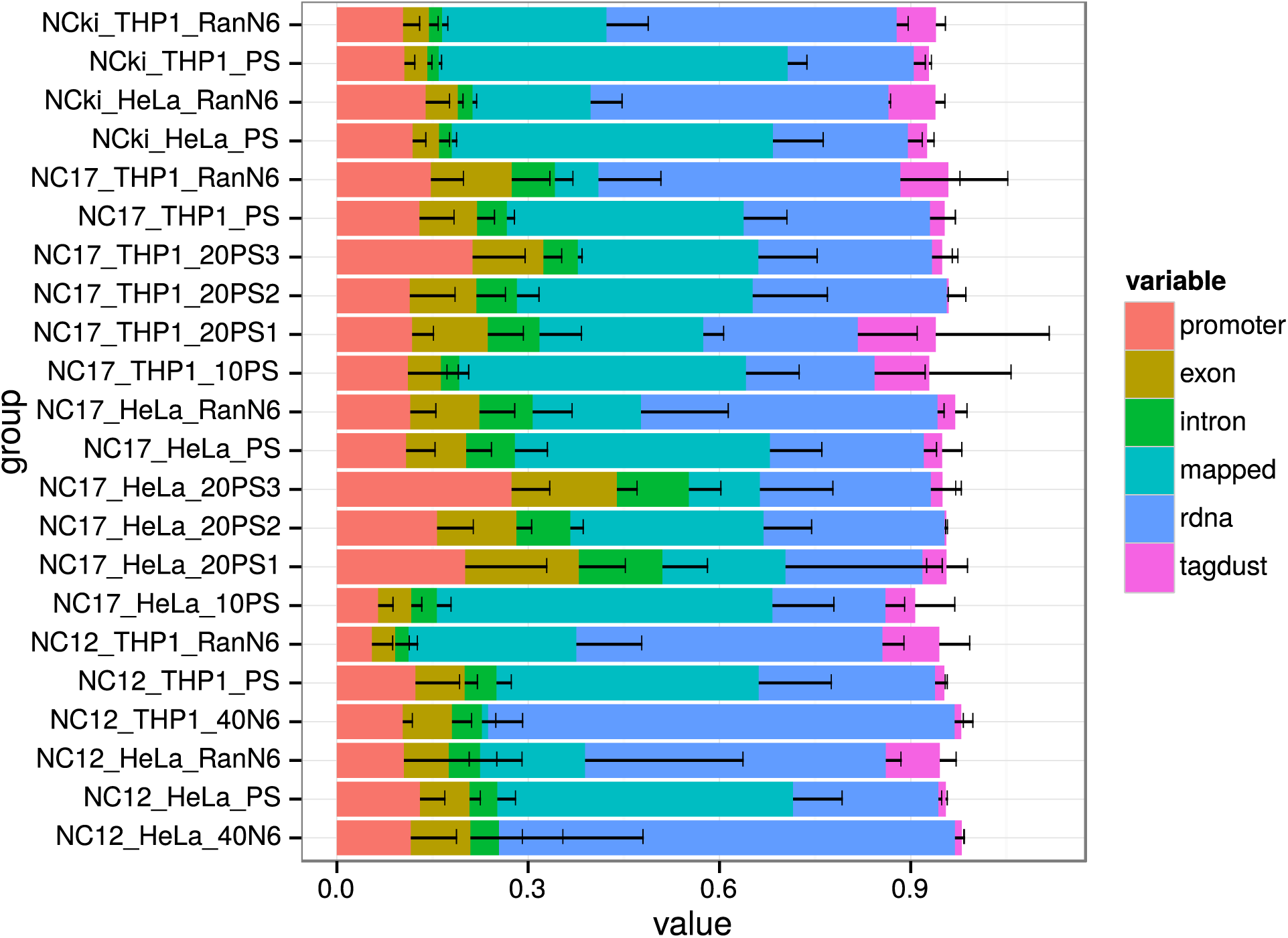

**Figure S2.**
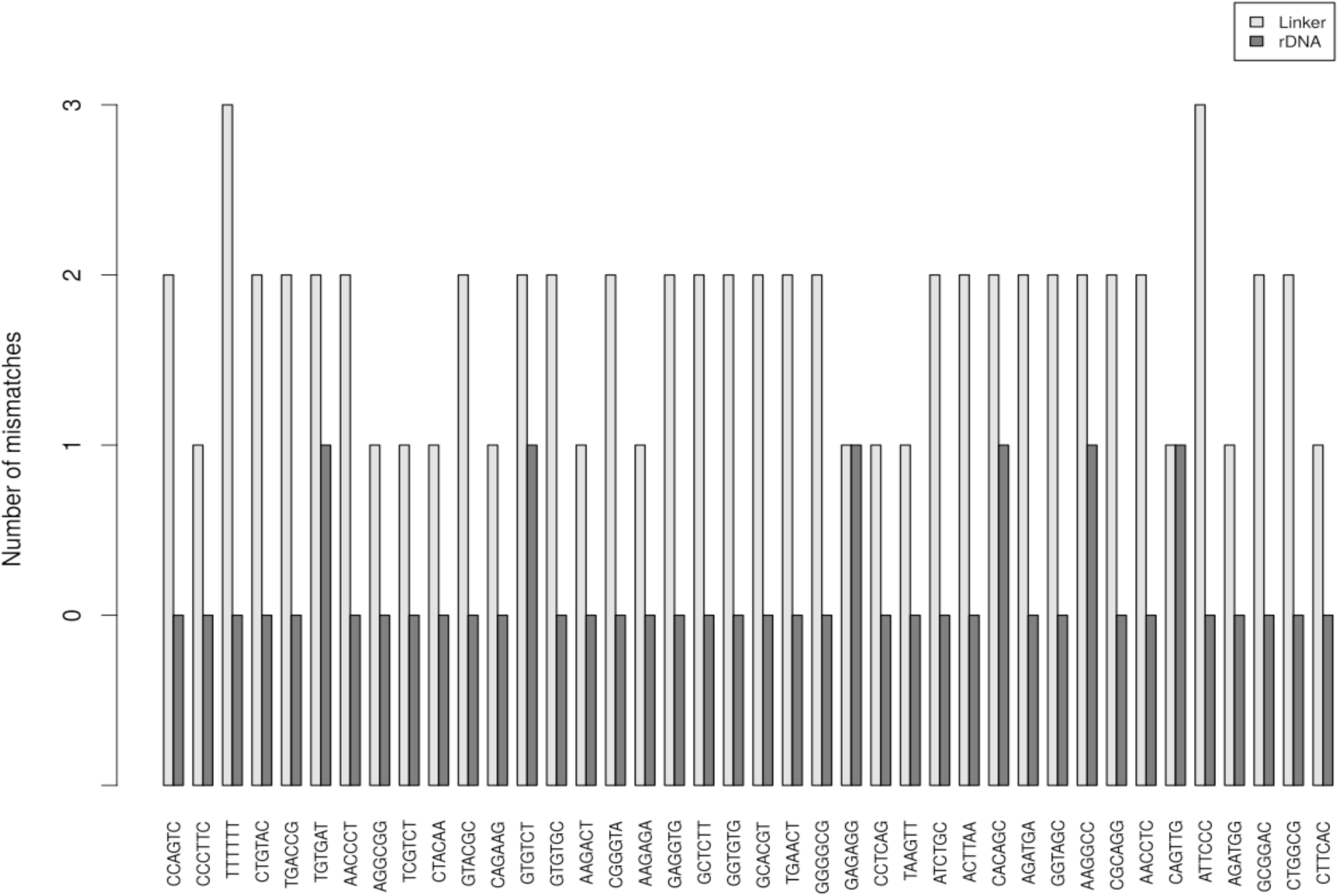

**Table S2.**

| 20PS set1 | 20PS set2 | 20PS set3 | 10PS |
| --- | --- | --- | --- |
| GCCAAA | CTGGCC | GCCAAA | GCCAAA |
| AGCAAA | TGTGCC | AAACAA | AAACAA |
| AAACAA | ATTGCC | TGCCAA | TGCCAA |
| ACACAA | CTACGC | CACACA | CACACA |
| TGCCAA | TATGGC | GTCACA | GTCACA |
| CAAACA | TTGTGC | GTGGCA | GTGGCA |
| CACACA | ACCACG | ATTTTA | ATTTTA |
| TGCACA | CACAGG | CACAAC | CACAAC |
| GTCACA | ACTGTG | AACCAC | AACCAC |
| TAGCCA | TGCCAT | TACCCC | TACCCC |
| GTGGCA | TGGCAT | CTGGCC |  |
| TGTTTA | GTGCAT | ATTGCC |  |
| ATTTTA | TTGTAT | TATGGC |  |
| CAAAAC | ATTTAT | ACCACG |  |
| CACAAC | TTTTAT | ACTGTG |  |
| GCTAAC | TGGCGT | TGGCAT |  |
| AACCAC | TGTTGT | TTGTAT |  |
| CTACCC | ATTTGT | TTTTAT |  |
| TACCCC | TTGCTT | TGTTGT |  |
| CTAGCC | TGTCTT | TTGCTT |  |

